# Atypical contribution of caspase-3 to melanoma cancer cell motility by regulation of coronin 1B activity

**DOI:** 10.1101/2024.06.27.601010

**Authors:** Kevin Berthenet, Serena Diazzi, Catherine Jamard, Kinga Stopa, Stefan Dragan, Deborah Fanfone, Trang Nguyen, Nathalie Al, Francois Virard, Olivier Meurette, Nikolay Popgeorgiev, Hector Hernandez-Vargas, Julien Ablain, Gabriel Ichim

**Author notes:** co-senior authors (Kevin Berthenet, Gabriel Ichim).

## Abstract

Recent studies have unveiled unexpected connections between cell death and cell motility. While traditionally recognized for their pro-apoptotic roles, caspases have emerged as regulators of physiological processes beyond cell death, including cellular differentiation and motility. In some particularly aggressive cancers like melanoma, caspase-3, a prominent executioner caspase, is unexpectedly and inexplicably highly expressed. Here, we describe a novel non-apoptotic role for caspase-3 in melanoma cell motility. Through comprehensive molecular and cellular analyses, we demonstrate that caspase-3 is constitutively associated with the cytoskeleton and crucially regulates melanoma cell migration and invasion *in vitro* and *in vivo*. Mechanistically, caspase-3 interacts with and modulates the activity of coronin 1B, a key regulator of actin polymerization, thereby promoting melanoma cell motility, independently of its apoptotic protease function. Furthermore, we identify specificity protein 1 (SP1) as a transcriptional regulator of *CASP3* expression, and show that its inhibition reduces caspase-3 expression and impairs melanoma cell migration. Overall, this study provides insights into the multifaceted roles of caspase-3 in cancer progression, highlighting its relevance as a novel target for anti-metastatic therapies.

## Introduction

Cell death and motility are two major physiological processes in living organisms, traditionally viewed as mutually exclusive phenomena. Apoptosis, which is the most prominent type of regulated cell death, is executed by lethal activation of cysteine-dependent aspartate-specific proteases called caspases ^1^. This occurs either through death receptor activation or, more commonly, as a consequence of mitochondrial outer-membrane permeabilization (MOMP) ^2^ ^3^. Caspases are expressed as inactive zymogens, also called pro-caspases ^1^. Executioner pro-caspase-3 and −7 are present as dimers and the cleavage within the inter-subunit linker trigger their apoptotic activation ^4^. Once fully active, executioner caspases, in particular caspase-3, cleave numerous protein substrates ensuring an efficient execution of apoptosis. Protein cleavage by caspases is also essential for the production and exposure of the so-called “find-me” and “eat-me” immunogenic signals that are involved in apoptotic cell recognition and clearance by macrophages ^5^ ^6^. Interestingly, chemical or genetic inhibition of caspase activation triggers a highly immunogenic form of cell death called caspase-independent cell death ^7^ ^8^.

One would expect cancer cells to have developed mechanisms to reduce or even eliminate caspase-3 expression, in order to dampen apoptosis and thus gain an even greater survival advantage. However, some aggressive cancers such as colon cancer and melanoma express high levels of caspase-3 ^9^ ^10^ ^11^ ^12^. Furthermore, the expression of certain caspases is associated with a poor prognosis in several cancers ^13,14^, and the therapeutic strategies based solely on lethal caspase activation are not as efficient as expected ^15^ ^3^.

Cancer cell motility is a hallmark of metastasis, which involves dynamic cell-intrinsic changes in cytoskeletal organization, focal adhesions and cell-to-cell contacts that ultimately extend to the tumor microenvironment, leading to modifications of intercellular matrix and cell invasion of adjacent tissue ^16^ ^17^. Cell motility is a manifestation of continuous processes of actin polymerization, cell adhesion and acto-myosin contractibility. Mechanistically, activation of the small GTPases Rac and Cdc42 promote actin-related protein 2 and 3 (ARP2/3)- dependent actin polymerization into F-actin in the motile cancer cells. This generates pro-migratory structures at the leading edge of invading cancer cells, such as lamellipodia or filopodia which are stabilized by cortical F-actin^17^. Metastatic melanoma is an ideal model for studying cancer cell motility since it is one of the most aggressive and deadliest form of cancer ^18^.

Surprisingly, caspases were described in several studies to participate in pathophysiological cell motility. For instance, *CASP3* knockout (KO) in colorectal cancer cells reduced their epithelial-to-mesenchymal transition (EMT) ^19^, while initiator caspase-8 was shown to be essential for neuroblastoma cell migration, by promoting calpain cleavage-mediated turnover of focal adhesion components ^20^ ^21^ ^22^. Interestingly, the cell motility-related effects of caspase-8 are independent of its proteolytic activity ^22^. In addition, pro-inflammatory caspase-11 cooperates with actin interacting protein 1 (Aip1) and cofilin-1 to promote actin depolymerization and leukocyte migration during inflammation ^23^. Here, we investigated whether the high level of expression of pro-caspase-3 (hereafter called caspase-3) by metastatic melanoma cells was linked to their high motility and aggressiveness.

## Results

### Caspase-3 is highly expressed in melanoma cells and tumors

To better characterize and understand the non-apoptotic functions of caspase-3 in melanoma, we first compared its mRNA expression in healthy skin tissue to a wide variety of tissues and organs. We found that *CASP3* has a high expression in skin, independently of sun exposure (**Fig.1A**). Then, focusing on melanoma, an aggressive type of skin cancer, we assessed whether *CASP3*, a powerful pro-apoptotic mediator, was mutated in melanoma tumors, using the Catalogue of Somatic Mutations in Cancer (COSMIC). Compared with major oncogenes like *BRAF* and *NRAS,* classically displaying genetic alterations in more than 50 and 20% of melanoma patients, respectively, *CASP3* was mutated in only 2% of cases (**Fig.1B**). However, among various initiator and effector caspases ^24^, *CASP3* stood out as being highly expressed in 39 melanoma cell lines from the Cancer Cell Line Encyclopedia (CCLE), encompassing both primary and metastatic-derived cells (**Fig.1C**). Interestingly, *CASP3* expression is also clinically relevant, since it significantly differentiated primary from metastatic melanoma tumors in the Cancer Genome Atlas Program (TCGA) melanoma dataset (**Fig.1D**).

**Figure 1.**
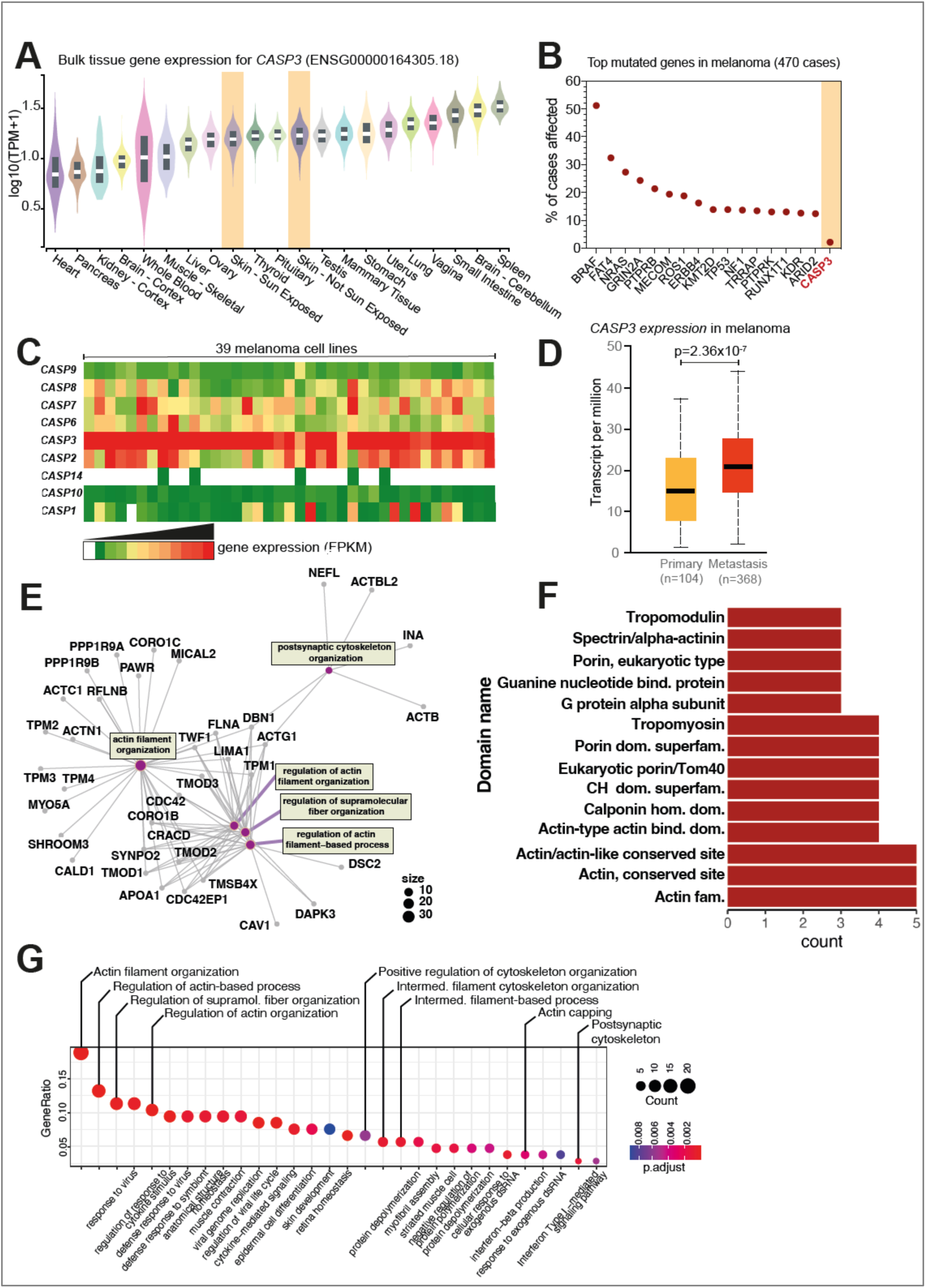
Caspase-3 is highly expressed in melanoma cells where it interacts with cytoskeletal proteins. **(A)** Analysis of *CASP3* expression in normal human tissues obtained from the GTEx (the Adult Genotype Tissue Expression) Portal. Skin tissue is highlighted. **(B)** Analysis of the most frequent mutations occurring in melanoma in the TCGA-SKCM dataset. *CASP3* mutation rate is highlighted in orange. **(C)** Analysis of caspase expression in 39 melanoma cell lines from the Cancer Cell Line Encyclopedia (CCLE). **(D)** Comparison of *CASP3* expression in primary (n = 104) and metastatic melanoma (n = 368) in the TCGA-SKCM dataset (p=2.36 x 10^-7^). **(E)** Clustering based on gene ontology (GO)-based classification of CASP3 interacting proteins identified by mass spectrometry after immunoprecipitation of CASP3-GFP (compared to GFP only precipitation) in WM852 cells. **(F)** Analysis of the type of protein domains contained in caspase-3 interacting partners in WM852. **(G)** Clustering based on the GO-based classification of CASP3-interacting proteins identified after immunoprecipitation of CASP3-GFP in WM793 and WM852 cells.

Next, to gain insight into the molecular and cellular pathways that might be regulated by caspase-3, *CASP3* expression was reduced using RNA interference in the human melanoma cell line WM793 (**Supp. Fig.1A**). This was instrumental for defining a gene signature comprising 310 differentially-expressed genes (DEGs) (both up- and downregulated) to discriminate between *CASP3*-depleted and control WM793 melanoma cells (**Supp. Fig.1B**). Melanoma patients with tumors harboring a strong association with genes that were upregulated following *CASP3* depletion (high UP signature) had a better prognosis (**Supp. Fig.1C, D**). Given these data and that most melanoma cells have maintained high levels of *CASP3* expression with very few mutations, its expression must confer melanoma cells with certain advantages, likely unrelated to the role of caspase-3 in apoptosis.

### Caspase-3 interacts with the cellular cytoskeleton in melanoma cells

To investigate the role of caspase-3 in melanoma cells, we followed two complementary approaches to fully characterize its interactome. We first stably expressed GFP or caspase-3-GFP fusion proteins in two different melanoma cell lines (WM793 and WM852) (**Supp. Fig.1E**). Using anti-GFP nanobodies coupled to magnetic agarose beads, we then immunoprecipitated caspase-3-GFP protein complexes and performed mass spectrometry analyses (**Supp. Fig.1 E-G**). The gene ontology (GO)-based classification of potentially interacting proteins for caspase-3 in WM852 cells revealed several GO clusters related to actin filament and cytoskeletal organization (**Fig. 1E**). In addition, an analysis of the type of protein domains contained in caspase-3-interacting partners in WM852, revealed an enrichment in actin-binding domains (**Fig. 1F**). To overcome possible limitations in this interactome analysis imposed by the choice of a single cell line, we clustered together the caspase-3-interacting proteins of WM852 and WM793 cells, and the GO analysis revealed a significant enrichment in GO terms related to “actin filament organization”, “regulation of actin-based processes” and “positive regulation of cytoskeleton organization” (**Fig. 1G**).

In a complementary approach designed to catch transient and weak protein-protein interactions that could have physiological relevance, we employed a proximity-dependent biotin identification method using a smal biotin ligase called BioID2 ^25^ ^26^. BioID2 was fused to either the N- or C-terminus of caspase-3, and the fusion proteins were expressed in two different cell lines (WM793 and WM852) in a doxycycline-inducible manner. Following biotin treatment, the biotinylated proteins were then immunoprecipitated with streptavidin-coated magnetic beads (**Supp. Fig.1H-L**). This gave us a comprehensive view of putative interacting proteins for caspase-3 in melanoma cells. Taken together, our data suggest that caspase-3 interacts with proteins involved in cytoskeletal organization.

### Caspase-3 participates in cytoskeletal organization in melanoma cells

Next, we further explored the role of caspase-3 in cytoskeletal organization by first performing an immunostaining of caspase-3 in WM852 cells. This revealed that a fraction of caspase-3 colocalizes with F-actin, primarily at the level of the cellular cortex (**Fig.2A**). Alternatively, melanoma cells expressing BioID2-caspase-3 were immune-stained with streptavidin-FITC, which revealed that biotinylation predominantly occurred in lamellipodia (**Fig.2B**). These results suggest that putative caspase-3-interacting partners are also present in lamellipodia. We confirmed these findings in cells expressing caspase-3 fused to BioID2 in the N-terminus (**Supp. Fig.2A**). In addition, sub-cellular fractionation experiments showed that, while most caspase-3 protein is cytosolic, a proportion is associated with the cytoskeletal fraction (**Fig.2C**). This was not the case for the executioner caspase-7 (**Fig.2C**).

**Figure 2.**
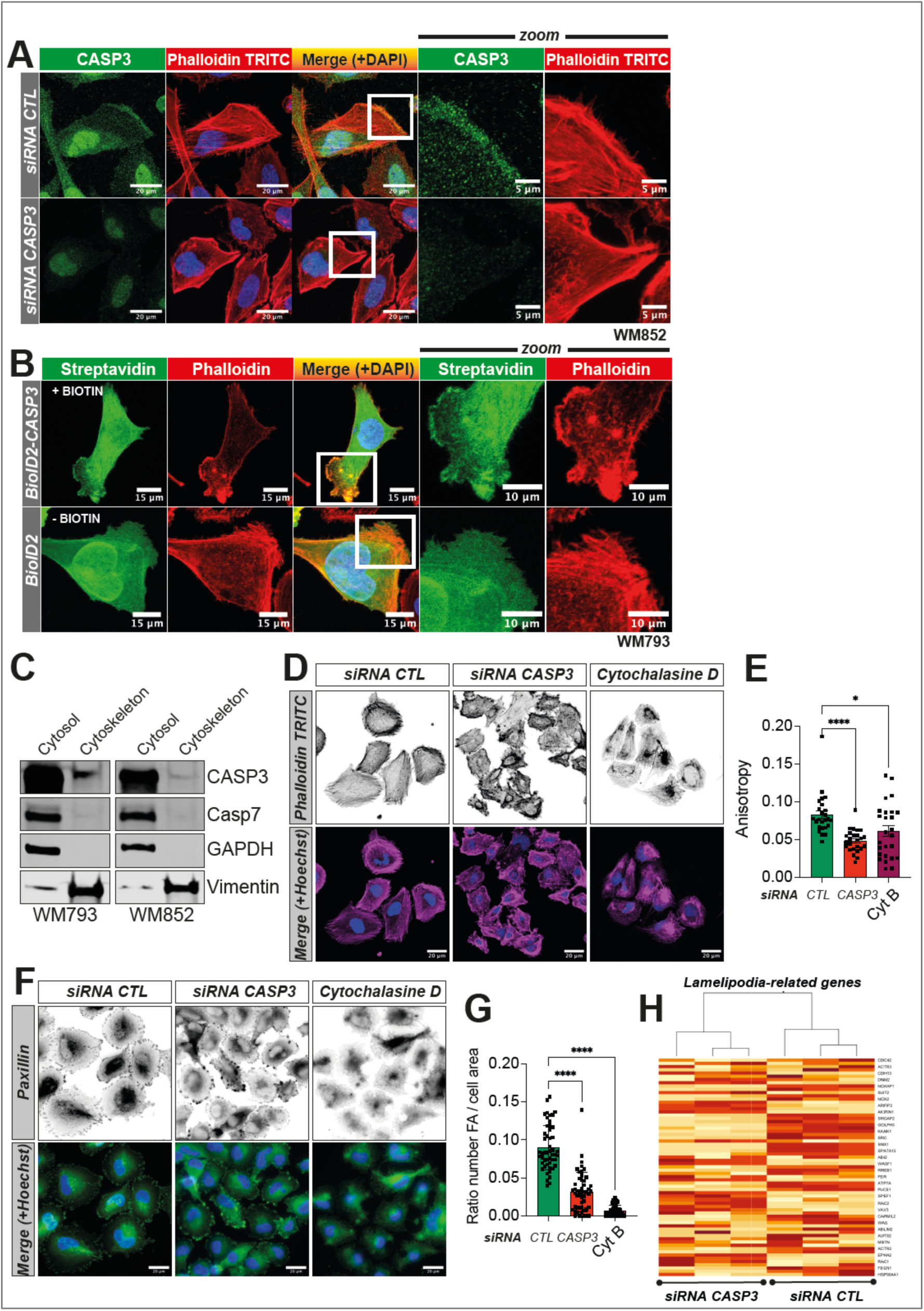
A fraction of Caspase-3 colocalizes with the cytoskeleton, specifically at the level of lamellipodia. **(A)** CASP3 and F-actin immunostaining in parental and CASP3-deficient WM852 cells. **(B)** Immunostaining using streptavidin-FITC of biotinylated proteins in WM793 cells expressing either BioID2 or BioID2-CASP3. **(C)** Immunoblotting for CASP3, CASP7, GAPDH and vimentin in cytosolic and cytoskeletal fractions of WM793 and WM852 cell lines. **(D)** F-actin staining in parental and CASP3-deficient WM793 cells. Cytochalasin D (1µM, 60 min) is an inhibitor of actin polymerization. **(E)** Quantification of F-actin anisotropy in parental and CASP3-deficient WM793 cells. **(F)** Paxillin immunostaining in parental and CASP3-depleted WM793. **(G)** Quantification of focal adhesions in parental and CASP3-depleted WM793 cells. **(H)** Clustering of lamellipodia-related genes from the GO-term GO:0030032 based on the RNA-seq perfomed in WM793 cells (siRNA control vs. siRNA CASP3).

To ascertain that caspase-3 is instrumental in cytoskeletal organization in melanoma cells, we down-regulated *CASP3* expression, and observed a complete disorganization of F-actin fibers, at a comparable level to cytochalasin B treatment, a known inhibitor of actin polymerization (**Fig.2D**). The anisotropy of F-actin fibers, which is their parallel alignment, is dramatically decreased by both *CASP3* down-regulation and cytochalasin B treatment (**Fig.2E**) ^27^. Since the architecture and functionality of focal adhesions are interconnected to maintain cytoskeletal organization, we tested whether caspase-3 downregulation impacted focal adhesions. Their number was lower in melanoma cells following acute reduction of caspase-3, as evidenced by paxillin staining, suggesting that melanoma cells lacking caspase-3 might have impaired cell to matrix adhesion and thus cell migration capacity (**Fig.2F, G**). Moreover, genes involved in regulating lamellipodia function, discriminated between control and caspase-3 downregulated melanoma cells (**Fig.2H**). Our data thus suggest that caspase-3 is associated with cytoskeletal proteins in melanoma cells.

### Caspase-3 is required for efficient melanoma cancer cell motility both *in vitro* and *in vivo*

We next sought to determine whether caspase-3 is instrumental for melanoma cell migration and invasion. Given that caspase-3-deficient melanoma cells displayed fewer focal adhesion points, we wondered whether this affected their level of adhesion to a matrigel-coated substrate. Caspase-3 knockdown in melanoma cells clearly impaired cell adhesion **(Fig.3A)**. Cellular tomography additionally showed that while control cells were completely attached, flat and able to expand lamellipodia, caspase-3 deficient cells were unable to efficiently attach and polarize (**Fig.3B**). Next, we conducted a series of IncuCyte live cell imaging-based cell migration and invasion assays, and established that reducing caspase-3 expression inhibited migration and invasion of WM793 and WM852 melanoma cells (**Fig.3C-H**). These findings were also complemented by chemotaxis assays, in which caspase-3 depleted cells displayed an impaired chemotaxis, i.e., cell motility (**Fig.3I-K**). This was equally valid for different siRNA and several *CASP3* KO cells generated through CRISPR/Cas9 gene editing (**Supp. Fig.3A-H**). Interestingly, this effect was specific for caspase-3 since knock-down of caspase-7 had no impact on melanoma cell migration (**Fig.3L**).

**Figure 3.**
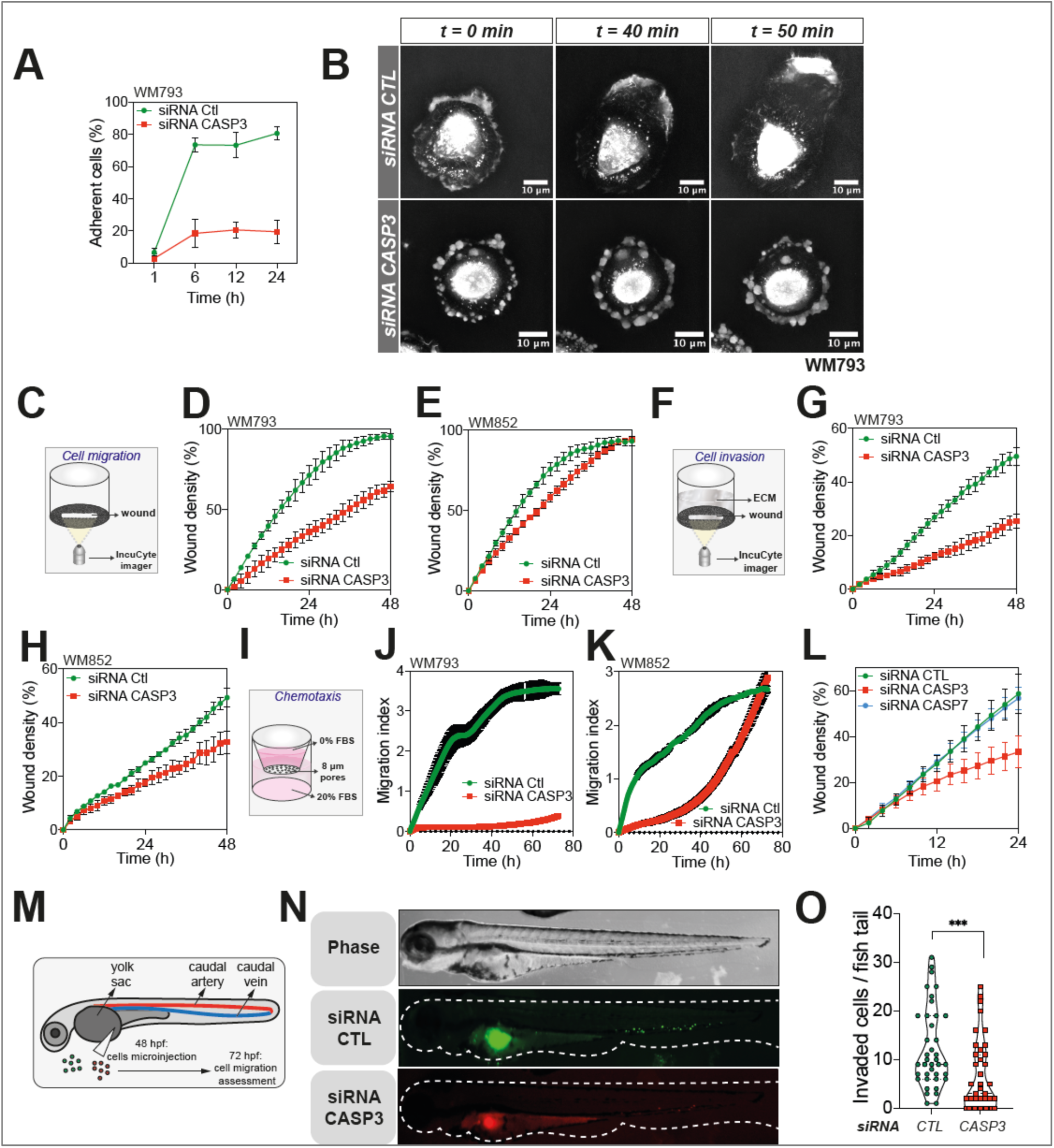
Caspase-3 controls melanoma cell motility. **(A)** Quantification of cell adhesion potential of parental and CASP3-depleted WM793 cells. **(B)** Time-lapse cellular tomography in WM793 cells. **(C)** Schematic representation of the wound healing assay using the IncuCyte live cell imager, which assesses the migratory ability of melanoma cells to close a wound. **(D-E)** Quantification of migration potential in parental and CASP3-deficient WM793 (**D**) and WM852 cells (**E**) through kinetic wound area measurement (data represent mean with SD of a representative experiment). **(F)** Schematic representation of the invasion assay through matrigel. **(G-H)** Measurement of the invasion potential of WM793 (**G**) and WM852 (**H**) depleted or not for CASP3 (data represent mean with SD of a representative experiment). **(I)** Schematic representation of the chemotaxis assay. **(J-K)** Measurement of control and CASP3-deficient WM793 (**J**) and WM852 cells (**K**) chemotaxis potential (data represent mean with SD of a representative experiment). **(L)** Analysis of the migration potential of control, CASP3 and CASP7 deficient WM793 cells through wound area measurement (data represent mean with SD of a representative experiment). **(M)** The zebrafish model of cancer metastasis. **(N)** DiD-labelled CASP3 deficient and DiO-labelled control WM793 cells were pre-mixed in equal numbers and injected in the zebrafish embryos yolk sac as shown in (**M**). A representative epifluorescence image of a whole embryo shows perivitelline homing and caudal blood vessels invasion of cancer cells. **(O)** Quantification of invaded metastatic cells per zebrafish embryo, (data represent mean with SEM, n=39 embryos from three independent experiments, t-test, *** p < 0.001).

To mirror these *in vitro* results in an *in vivo* setting, we used a zebrafish (*Danio rerio*) model to test early metastatic dissemination. The zebrafish provides a highly versatile system to evaluate cancer cell migration and invasion within a functional circulatory system, and the transparency of the early-stage embryos facilitates quantitative evaluations ^28^ ^29^. Parental and caspase-3-deficient cells were stained with the vital fluorescent dyes DiI and DiD, respectively, and microinjected into the perivitelline cavity of 48-h-old post fecundation (hpf) zebrafish embryos (**Fig.3M**). Twenty-four hours later the metastatic dissemination of melanoma cells within the tail vascular network was scored, and confirmed that caspase-3 depletion impaired metastatic dissemination *in vivo* (**Fig.3N, O**). Hence, our results propose caspase-3 as a regulator of melanoma cell motility and involved in metastatic dissemination.

### Caspase-3 drives melanoma cell motility independently of its apoptotic protease activity

Our data demonstrate that caspase-3 is involved in cancer cell motility, yet whether this occurs independently of its pro-apoptotic function is unknown. To address this, we first used the pan-caspase inhibitor Q-VD-OPh to block all residual caspases that might be activated at non-lethal levels. As depicted in **Supp. Fig.3I**, chemical caspase inhibition had no effect on melanoma cell motility. To strengthen this finding, we also used a BiFC (bi-molecular fluorescence complementation)-based reporter, highly sensitive to both non-apoptotic and apoptotic executioner caspase activation ^30^ ^31^ ^32^. While a classical apoptotic stimulus such as actinomycin D triggers massive caspase activation, melanoma cells had no residual caspase activity that could participate in cellular motility (**Supp. Fig.3J, K**). To completely block non-lethal caspase activation via the intrinsic mitochondrial pathway, we also used CRISPR/Cas9-mediated knock-out of both pore-forming proteins BAX and BAK (**Supp. Fig.3L**). Melanoma cells lacking both BAX and BAK, had the same migratory capacity as parental cells (**Supp. Fig.3M, N**), confirming our previous results. Our data thus support an effector role of caspase-3 in promoting melanoma cell motility, independently of its well-established apoptotic protease function.

### Caspase-3 facilitates melanoma metastasis *in vivo*

To enhance our data on the non-canonical function of caspase-3 in promoting melanoma dissemination, we next took advantage of a model of genetically-engineered primary melanoma in adult zebrafish, where transgenesis is efficient and rapid. More precisely, a MiniCoopR vector for melanocyte-specific overexpression of human *CASP3* was microinjected in zebrafish embryos that are p53 deficient and express BRAF V600E ^33,34^. Two to six months later, primary melanoma tumors were observed (**Fig.4A**). Figure 4B, demonstrates the efficacy of caspase-3 overexpression in primary zebrafish melanoma, compared to GFP used as an overexpression control. Importantly, caspase-3 overexpression did not alter the growth of the primary tumor, neither decreasing it through excessive cell death induction nor enhancing it, since the tumor-free survival was comparable between fish bearing GFP and caspase-3-expressing tumors (**Fig.4C**). Next, pigmented primary tumors overexpressing either GFP or caspase-3 were dissociated and transplanted subcutaneously into the dorsal cavity of secondary transparent Casper zebrafish. Three weeks later, tumor dissemination was evaluated by quantifying the patches of melanoma cells that disseminated beyond the midline (**Fig.4D**). This revealed that melanoma cells overexpressing caspase-3 have a significant gain of metastatic potential *in vivo* (**Fig.4E, F**).

**Figure 4:**
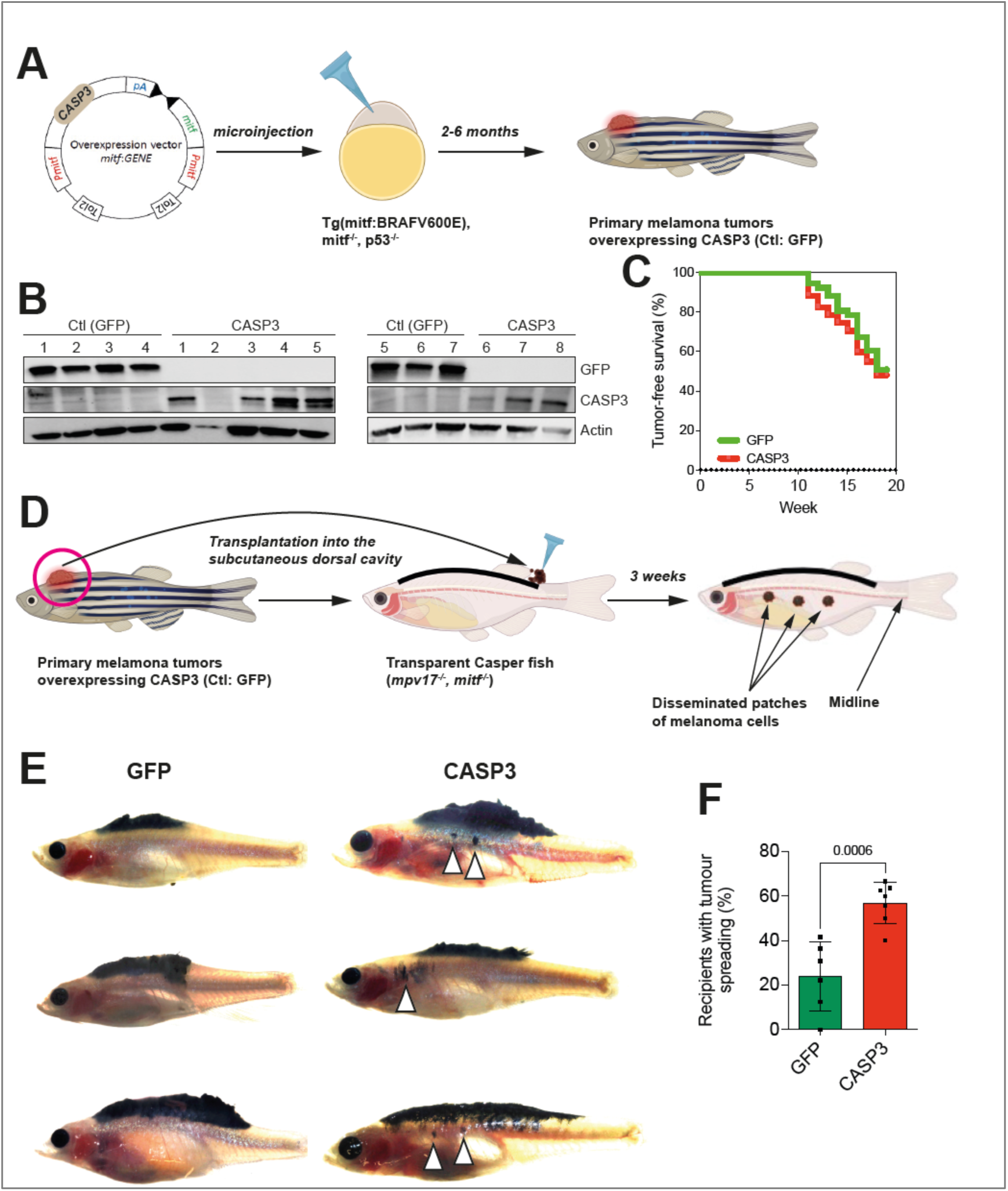
Caspase-3 facilitates melanoma metastasis in vivo. **(A)** Workflow for the generation of primary zebrafish melanomas upon microinjection of the transposon-based MiniCoopR vector into one-cell stage embryos. **(B)** Western blot analysis showing the expression levels of Caspase-3 (CASP3) and Green Fluorescent Protein (GFP/Ctl) in primary zebrafish melanoma tumors. **(C)** Tumor-free survival curves of Tg (mitfa:BRAF^V600E^), tp53^−/−^, mitfa^−/−^ zebrafish injected with vectors for the overexpression of CASP3 or GFP (Ctl). Statistical analysis was performed using the log-rank test. **(D)** Workflow for the transplantation of primary zebrafish melanomas into the dorsal cavity of secondary recipient Casper fish. **(E)** Representative images of Casper fish transplanted with primary zebrafish melanomas overexpressing CASP3 or GFP. Arrows indicate the location of disseminated melanoma cells. **(F)** Quantification of the percentage of secondary recipients with tumor spreading, comparing control GFP (n = 6) and CASP3 (n = 7) overexpressing tumors. Data are presented as mean ± standard deviation (S.D.); statistical analysis was performed using two-tailed t-test.

### Efficient melanoma cell migration requires caspase-3-mediated coronin 1B activation

To gain further insight into the mechanisms regulating melanoma cell migration by caspase-3 expression, we drew the list of putative caspase-3-interacting partners in WM793 cells, combining caspase-3-GFP IP-MS and BioID2 proteomic approaches. One of the hits we obtained was coronin 1B. For further mechanistic studies we focused on this candidate since it is a protein mediating cell motility by inhibiting the actin-nucleating activity of the Arp2/3 complex ^33^ (**Fig.5A, Supp. Fig.5A**). Importantly, proteomic data were first confirmed by co-immunoprecipitation experiments in WM793 and WM852 cells, where coronin 1B specifically interacted with caspase-3-GFP (**Fig.5B, Supp. Fig.5B**). In a complementary approach, we used proximity ligation assay (PLA), which revealed a specific interaction between the two proteins (**Fig.5C, D and Supp. 5C, D**). Next, we immune-stained both endogenous caspase-3 and coronin-1B in WM793 cells. As shown in **Fig. 5E**, coronin 1B localizes at the leading edge of melanoma cells, co-localizing with a fraction of caspase-3 and F-actin, while the co-localization between coronin 1B and F-actin is lost when caspase-3 in downregulated (**Fig.5E**, line scans). In line with its role in cell motility, downregulation of coronin 1B with two different siRNAs has an inhibitory effect on melanoma cell migration (**Fig.5F**).

**Figure 5:**
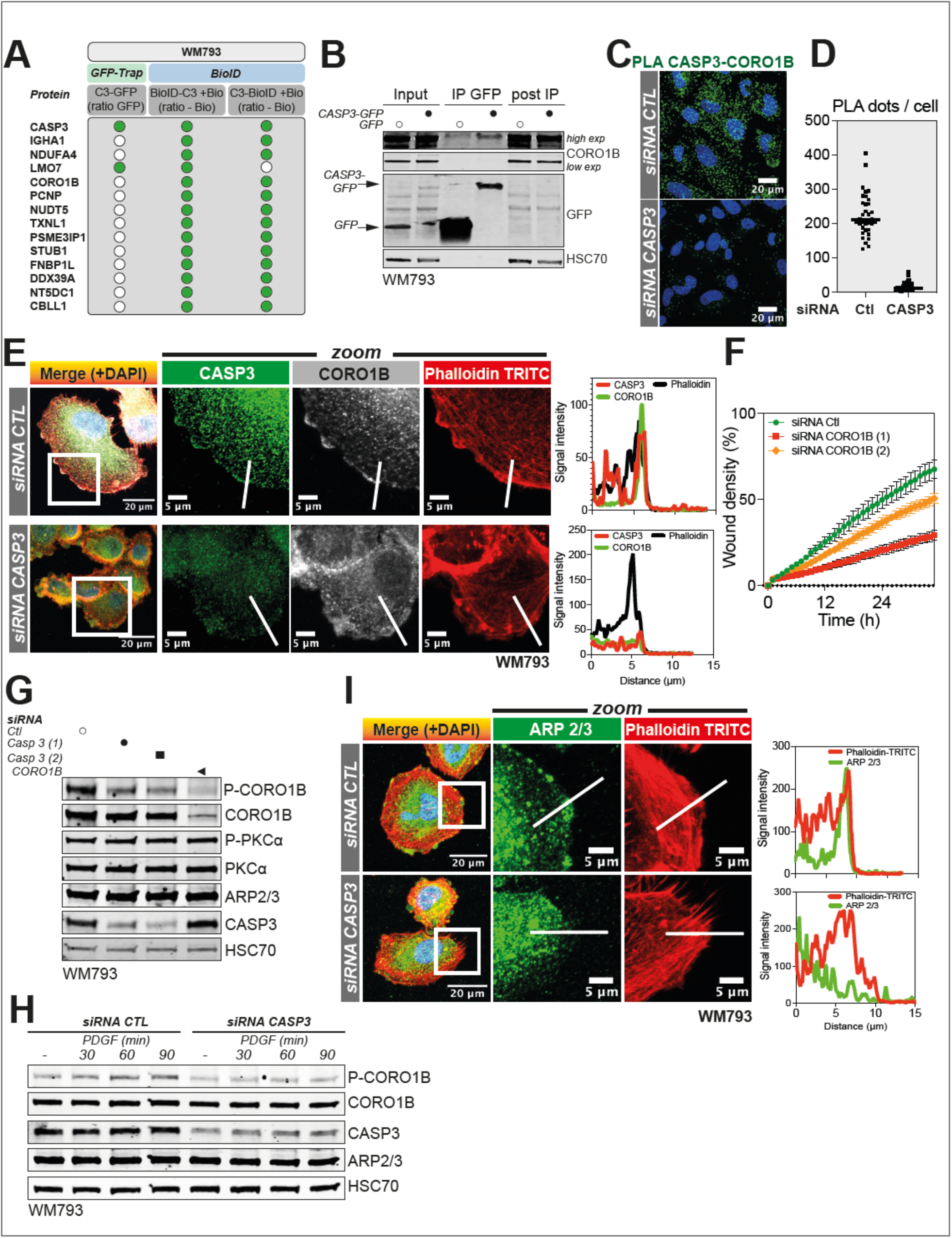
Caspase-3 sustains melanoma cell motility by regulating CORO1B activity. **(A)** Summary table with the most frequent CASP3 putative interacting protein partners identified following CASP3-GFP pulldown and proximity biotinylation in melanoma cells expressing CASP3 fused with BioID2, in either Nter or Cter. **(B)** Analysis of CASP3-GFP and CORO1B interaction after immunoprecipitation (IP) of GFP protein complexes in GFP- or CASP3-GFP in WM793 melanoma cells. **(C)** Analysis of the proximity between endogenous CASP3 and CORO1B proteins in control and CASP3-deficient WM793 using a Proximity Ligation Assay (PLA). **(D)** Quantification of PLA signal in control (n = 35 cells) and CASP3 (n = 31 cells) deficient WM793 cells. **(E)** Left panel: Analysis of CASP3, CORO1B and F-actin localization by immunofluorescence in control and CASP3-deficient WM793 cells. Right panel: Signal intensity measurement of CASP3, CORO1B and F-actin along the indicated corresponding line in control and CASP3-deficient WM793 in the left panel. **(F)** Measurement of cell invasion through Matrigel in control and CORO1B-deficient WM793 (data represent mean with SD of a representative experiment). **(G)** Analysis by immunoblotting of P-CORO1B, CORO1B, P-PKCα, PKCα, CASP3 in WM793 transfected with two different siRNA for CASP3 and CORO1B. HSC70 serves as a loading control. **(H)** Analysis by immunoblotting of P-CORO1B, CORO1B, ARP2/3 and CASP3 in control and CASP3-deficient WM793 cells that were serum-starved for 24 h and treated with PDGF (20 ng/mL) for the indicated time. HSC70 serves as a loading control. **(I)** Left panel: Analysis of ARP2/3 and F-actin localization by immunofluorescence in control and CASP3-deficient WM793 cells. Right panel: Signal intensity measurement of ARP2/3 and F-actin along the indicated corresponding line in control and CASP3-deficient WM793 in the left panel.

Coronin 1B was shown to be activated through phosphorylation of the serine-2 residue (Ser-2) by protein kinase C (PKC) promoting cell motility ^34^. Interestingly, we found at steady-state, that reducing caspase-3 expression lowers coronin 1B phosphorylation (**Fig.5G and Supp. Fig.5E**). This was confirmed by immunofluorescence in caspase-3-depleted cells, in which phospho-coronin 1B was lost in the lamellipodia (**Supp. Fig.5F**). We then assessed whether known PKC activators (ester phorbol 12-myristate 12-acetate or PMA and PDGF) could efficiently activate coronin 1B in parental and caspase-3 depleted melanoma cells ^35^ ^36^. As shown in **Fig.5H** and **Supp. Fig.5G, H**, both PMA- and PDGF-induced coronin 1B phosphorylation are delayed by caspase-3 depletion. This was mirrored by a defective ARP2/3 complex recruitment to the F-actin rich structures, such as the cellular leading edge (**Fig.5I**). Of note, caspase-3 depletion did not affect coronin 1B protein stability, as assessed by a chase assay using the protein translation inhibitor cycloheximide (**Supp. Fig.5I**). Altogether, we demonstrate here that caspase-3 has a non-canonical role in melanoma cell motility by regulating coronin 1B activation and its subcellular localization.

### Caspase-3-mediated cancer cell motility requires an intact SP1 transcriptional regulation

Since melanoma cell motility underlies its aggressiveness, we next sought to determine the regulatory mechanisms underlying high caspase-3 expression, in order to target this mechanism to reduce cell migration and invasion. We focused on Specificity Protein 1 (SP1) based on a previous study showing that the *CASP3* promoter contains three binding motifs for this transcription factor (**Fig.6A**) ^37^. Interestingly, SP1 is known to be constitutively overexpressed in several cancers and associated with poor prognosis ^38^. Here, the mRNA expression of both *CASP3* and *SP1* were positively correlated in melanoma samples (**Fig.6B**). To test whether SP1 regulated *CASP3* expression in melanoma cells, we first used siRNA to deplete SP1 in WM793 and WM852 cells. This led to a marked decrease in caspase-3 transcript and protein expression (**Fig.6C, D, Supp. Fig.6A, B**). Conversely, when SP1 was overexpressed in WM852 cells, this caused higher levels of caspase-3 protein (**Fig.6E**). At the functional level, SP1 knockdown impaired melanoma cell migration and invasion both *in vitro* and in zebrafish larvae (**Fig.6F-H**). The SP1 inhibitor, mithramycin A, efficiently reduced *CASP3* mRNA and protein expression (**Supp. Fig.6C, D, Fig.6I**) ^39^ and consistently blocked melanoma cell migration both *in vitro* and *in vivo* (**Fig.6J-L**). Taken together, these data identify SP1 as an actionable regulator of caspase-3 expression that could be pharmacologically modulated to reduce the pro-migratory role of caspase-3 in melanoma.

**Figure 6:**
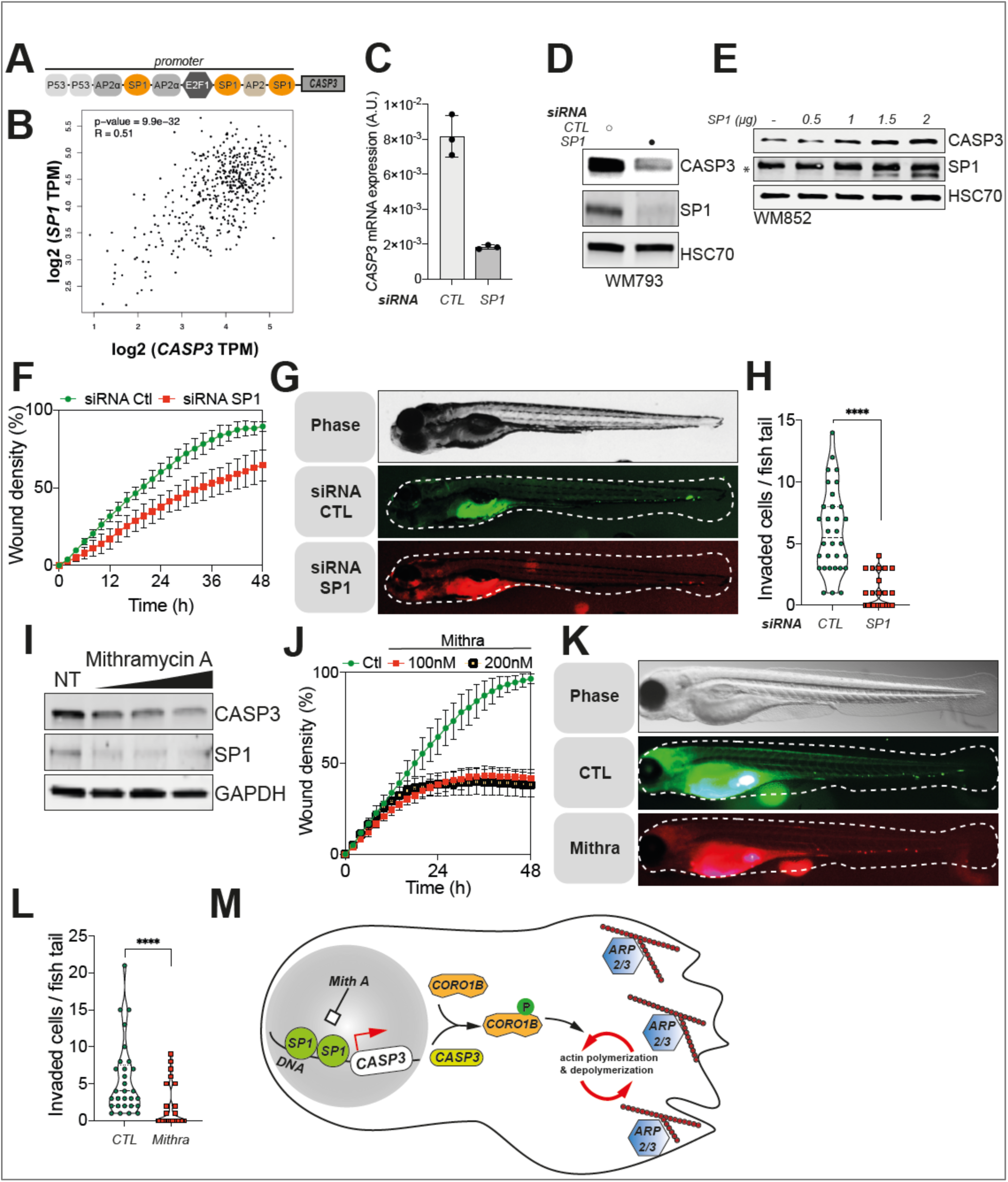
Reduction of CASP3 expression through SP1 inhibition limits melanoma dissemination potential. **(A)** Schematic representation of *CASP3* promoter described by Sudhakar *et al.*, 2008, FEBS J. **(B)** Analysis of *SP1* and *CASP3* mRNA expression in melanoma patients with melanoma (R = 0.51; p-value = 9,9 * 10^-32^). **(C)** *CASP3* mRNA expression in control and SP1-deficient WM793 cells, relative to GAPDH. **(D)** Western blot analysis of CASP3 and SP1 in control and SP1 siRNA transfected WM793 cells. HSC70 serves as loading control. **(E)** Analysis of CASP3 expression in WM852 transfected with increasing concentration of a SP1-expressing plasmid. HSC70 serves as a loading control. **(F)** Quantification of the migration potential of parental and SP1-deficient WM793 cells through wound area measurement (data represent mean with SD of a representative experiment). **(G)** DiD-labelled SP1-deficient and DiO-labeled control WM793 cells were pre-mixed in equal numbers and injected as shown in **3M**. A representative epifluorescence image of a whole embryo shows perivitelline homing and caudal blood vessels invasion of cancer cells. **(H)** Quantification of invaded metastatic cells per embryo tail veins (data represent mean with SEM, n=32 embryos from three independent experiments, t-test, **** p < 0.0001). **(I)** Analysis of CASP3 and SP1 expression in WM793 treated with increasing doses of mithramycin A (100 nM, 200 nM and 300 nM). GAPDH serves as a loading control. **(J)** Wound healing migration assay for WM793 cells treated with 0, 100 or 200 nM of mithramycin A (data represent mean with SEM of a representative experiment). **(K)** DiD-labelled mithramycin A treated (100 nM) and DiO-labelled control WM793 cells were pre-mixed in equal numbers and injected as shown in **3M**. A representative epifluorescence image of a whole embryo shows perivitelline homing and caudal blood vessels invasion of cancer cells. **(L)** Quantification of invaded metastatic cells per embryo (data represent mean with SD, n=29 embryos from three independent experiments, t-test, **** p < 0.0001). **(M)** Model: Caspase-3 in highly expressed in melanoma cells, most likely through a transcriptional regulation by SP1, that could be therapeutically blocked by treatment with mithramycin A. Caspase-3 interacts with coronin 1B (CORO1B), promoting its phosphorylation and localization at the cellular leading edge. CORO1B then interacts with ARP2/3 complex at the leading edge contributing to continuous cycles of actin polymerization and depolymerization. Caspase-3 downregulation leads to lower CORO1B phosphorylation, defects in its localization at the leading edge of melanoma cells and therefore decreased cell motility.

## Discussion

Caspases, in particular caspase-3, are well-known executioners of apoptosis, and their expression is frequently aberrant or impaired in cancer cells. Here, focusing on melanoma cells that are both very aggressive and express high caspase-3 levels, we describe a previously unknown function of this protease in cancer cell motility. Indeed, using complementary proteomic methods, we show that caspase-3 interacts with several cytoskeletal proteins. When caspase-3 expression is disrupted, melanoma cells display a disorganized F-actin cytoskeleton and fewer focal adhesions. Consequently, caspase-3-deficient melanoma cells are less responsive to chemotactic cues, migrate and invade less, both *in vitro* and *in vivo*, when injected in the vasculature of zebrafish embryos. Importantly, this non-canonical role is independent of its role as an apoptotic protease. Additionally, our study identified the transcription factor SP1 as key regulator of caspase-3 expression in melanoma cells. When SP1 regulation is disrupted, caspase-3 protein levels decrease, negatively affecting cancer cell motility. Furthermore, caspase-3 interacts with coronin-1B, a key regulator of actin dynamics ^33^. This interaction is required for guiding coronin-1B to the leading edge of migrating cancer cells and activating it properly.

Specifically, caspase-3 modulates the activating phosphorylation of coronin-1B and the correct subcellular localization of the ARP2/3 complex (model in **Fig.6M**).

Although many studies have suggested that increased caspase-3 expression is associated with improved patient outcome, some studies reported a positive correlation between higher caspase-3 expression, disease progression and poor outcomes for cancer patients ^9^ ^10^ ^12^ ^14^ ^40^. The role of caspase-3 as an executioner protease during apoptosis, implies that higher caspase-3 expression is permissive to either lethal or non-lethal caspase activation. Despite the common understanding that apoptotic cells are signaling-inert, recent research suggests that apoptosis results in the secretion of various signaling molecules that can remodel the tumor microenvironment. More precisely, apoptotic cancer cells secrete prostaglandins, FGF2 or ATP, which can have pro-oncogenic effects, such as compensatory proliferation and enhanced survival, on the remaining cancer cells ^15,41–44^. When effector caspases are not fully activated to lethal levels, they could also enhance oncogenesis by inducing DNA damage, mutagenesis, activate JNK and thus contribute to enhanced cell aggressiveness ^3,31,45,46^. In this study we provide an alternative explanation on the link between an increased caspase-3 expression and a poor patient outcome in some cancers. Independently of its cysteine protease role, we show here that caspase-3 also promotes cancer cell motility, at least in melanoma cells. Other caspases, such as caspase-8 and −11 were reported to enhance cell motility, however whether this function is related to their cysteine protease activities remains to be addressed with unbiased proteomic assays to identify their interacting partners. Aside from caspases, other proteases contribute to controlling cell migration: (i) calpains are regulators of both integrin-mediated adhesion and actin polymerization at the leading edge of the cell, (ii) matrix metalloproteinases are essential for extracellular matrix degradation and therefore facilitate the invasion of tumor cells, (iii) while extracellular cysteine cathepsin proteases cleave cell-cell adhesion molecules, enhancing tumor cells invasiveness ^47,48^ ^49,50^ ^51^. Such proteins could also have protease-independent roles, which may constitute novel targets for highly invasive cancer cells.

In this study, we found that SP1 is a major regulator of caspase-3 expression, with a significant correlation between *SP1* and *CASP3* mRNA in melanoma tumors, supporting the idea that SP1 inhibition could be envisioned as a migrastatic therapeutic intervention. Although discontinued from manufacturing in 2000, the SP1 inhibitor mithramycin A (known by the trade name of mithracin) was shown to hinder the metastatic potential of lung cancer and salivary adenoid cystic carcinoma cells ^52^ ^53^. Whether this effect relies, at least partly, on the capacity of caspase-3 to control cell motility, requires further investigations. Our study uncovered a functional interaction between caspase-3 and coronin-1B, leading to the optimal activation of the later. This reveals intriguing perspectives about identifying the exact interacting domain(s) and interface(s) between the two proteins and whether designing small molecules to interfere with the binding might also be a promising cancer migrastatic intervention.

It is important to bear in mind the possible bias and limitations of our study. The novel non-canonical role of caspase-3 in cancer cell motility was tested exclusively in melanoma, and further investigations will be required to assess the generalization of these finding for other types of aggressive cancers such as pancreatic adenocarcinoma or glioblastoma. Ideally, this could also be assessed in genetically-engineered mouse models predisposed to spontaneous metastatic dissemination, where caspase-3 expression could be conditionally modulated. The main findings of this study could also be tested throughout metastatic progression, starting with intravasation, surviving the shear stress in the blood stream, extravasation and metastatic colonization.

In summary, we report here that caspase-3 has a non-apoptotic and protease-independent role in sustaining melanoma cell motility. This offers new perspectives for testing whether certain mechanistic findings of this study could contribute to various other roles for caspase-3 in cellular differentiation, neuronal function, wound repair or regeneration.

## Acknowledgements

This work was supported by funding from LabEx DEVweCAN (University of Lyon), the National Cancer Institute (INCa PLBIO21-003 to GI and PLBIO23-255 to JA), Centre National de la Recherche Scientifique and La Ligue Nationale Contre le Cancer (ATIP-Avenir 2019 to JA), Fondation de France (00108257 to KB and 00130886/WB-2022-41429 to SD), Fondation ARC pour la recherche sur le cancer (ARCPJA2022060005168 to JA) and the Polish National Agency for Academic Exchange (NAWA) (BPN/BEK/2023/1/00339/U/00001). We thank Brigitte Manship for reviewing the manuscript. We also thank Pierre Sujobert, Lorric Delage and Elodie Cascales for their contribution to the set-up of BioID2 and CRISPR/Cas9 gene editing via electroporation experiments. This study was also facilitated by the following CRCL core facilities: “Gilles Thomas” bioinformatics platform, Flow cytometry core facility (CYLE), Biological sample management platform (PGEB) and Cell imaging platform (PIC), as well as by the PRECI Aquatics facility (SFR Biosciences - UAR3444/US8). Finally, we acknowledge Adeline Page and Frédéric Delolme from the Protein Science Facility at the SFR Biosciences (UAR3444/CNRS, US8/Inserm, ENS de Lyon, UCBL) for the Mass spectrometry analyses.

## Authors’ contributions

<u>Conceptualization:</u> K. B., G. Ichim; <u>Methodology:</u> K. B., G. Ichim, J.A., F.V., O. M., N. P., H. H.-V.; <u>Formal</u> <u>analysis</u>: K. B, G. Ichim, S.D., F. V., O. M., H. H.-V.; <u>Investigation</u>: K. B., S. D., C. J., K. S., S. D., D. F., T. N., N. A., G. Ichim; <u>Resources</u>, J. A., G. Ichim; <u>Writing</u> – Original Draft and Editing, K. B. and G. Ichim.; All authors reviewed and edited the manuscript; <u>Supervision</u>, G. Ichim, K. B., J. A., O. M.; <u>Project administration and funding acquisition</u>, G. Ichim and K. B.

## Material and methods

### Cells and reagents

Melanoma cell lines (WM793, WM852, and A375) and 293T cells were cultured in Dulbecco’s Modified Eagle’s Medium (DMEM) supplemented with 10% fetal bovine serum (FBS) (Eurobio, CVFSVF00-01), 2 mM glutamine (Thermo Fisher Scientific, 25030-024), non-essential amino acids (Thermo Fisher Scientific, 11140-035), 1 mM sodium pyruvate (Thermo Fisher Scientific, 11360-039), and penicillin/streptomycin (Thermo Fisher Scientific, 15140-122). Cells were cultured in a humidified atmosphere at 37°C and routinely checked for mycoplasma contamination. When indicated, cells were treated with the following reagents: Cytochalasin D (Sigma-Aldrich, C8273-1MG), qVD-OPh (Clinisciences, HY-12305-5mg), biotin (Sigma-Aldrich, B4501-5G), actinomycin D (Sigma-Aldrich, 9415-5MG), ABT-263 (ApexBio, A3007) and mithramycin A (Sigma-Aldrich, 5303100001).

### Establishment of GFP or CASP3-GFP stable melanoma cells

Parental WM793 and WM852 melanoma cells were transfected with either p-EGFP-N3 or p- EGFP-N3-CASP3 plasmids using Lipofectamine 2000 (ThermoFisher Scientific, 11668019) according to the manufacturer’s protocol. Forty-eight hours after the transfection, cells were selected for stable overexpression of GFP or CASP3-GFP by incubation with 500 µg/mL neomycin (Sigma-Aldrich, G8168). Transfected cells were subsequently sorted using a FACS ARIA cell sorter (BD Biosciences) to isolate a homogeneous population exhibiting uniform GFP expression.

### Generation of VC3AI stable melanoma cells

293T cells (1.5 x 10^6^ cells in a 10-cm dish) were transfected with the pCDH-puro-CMV-VC3AI lentiviral vector (Addgene, 78907) using Lipofectamine 2000. Concurrently, the lentiviral packaging plasmids pVSVg (Addgene, 8454) and psPAX2 (Addgene, 12260) were co-transfected to facilitate virus production. After 48 h, virus-containing supernatants were collected, filtered, and used to infect WM793 and WM852 melanoma cells in the presence of 1 µg/mL polybrene (Sigma-Aldrich, H9268). Stably transduced cells were selected by culturing in 1 µg/mL puromycin (Invivogen, ant-pr-1) for two days post-infection.

### Establishment of stable melanoma cells for doxycycline-inducible expression of BioID2 and CASP3 fused with BioID2

WM793 and WM852 melanoma cell lines were transfected with pITR-mycBioID2, pITR-mycBioID2-CASP3, and pITR-CASP3-mycBioID2 constructs, using the Sleeping Beauty transposon system (Kowarz et al., 2015). Plasmids were co-transfected with the pCMV(CAT)T7-SB100 encoding SB100X transposase using Lipofectamine 2000. Transfected cells were selected with 100 µg/mL zeocin (Invivogen, ant-zn) to establish stable cell lines.

### Generation of *CASP3*, *BAX* and *BAK*-deficient melanoma cells using CRISPR/Cas9-gene editing

293T cells (1.5 x 10^6^ cells in a 10-cm dish) were transfected with lentiCRISPR v2 h*CASP3* (see plasmid construct part for design), lentiCRISPR v2 h*BAX* (Addgene, 129580) and/or LentiCRISPR v2 h*BAK1* (Addgene, 129579) plasmids using Lipofectamine 2000. Lentiviral packaging was facilitated by co-transfecting with pVSVg and psPAX2 plasmids. Two days post-transfection, virus-containing supernatants were collected, filtered, and used to infect WM793 and WM852 melanoma cells, in the presence of 1 µg/mL polybrene. Cells were selected for stable gene knockout by culturing in 10 µg/mL blasticidin (Invivogen, ant-bl).

### Generation of *CASP*3-Deficient Melanoma Cell Lines with CRISPR/Cas9 via electroporation

CASP3-deficient WM793 were established using the CRISPR genome editing methodology developed by IDT according to the manufacturer’s instructions. *CASP3*-targeting CRISPR-Cas9 crRNA was designed using the Alt-R™ CRISPR HDR Design Tool (https://www.idtdna.com/pages/tools/alt-r-crispr-hdr-design-tool) and its sequence is detailed in Table S1. The *CASP3*-targeting CRISPR-Cas9 crRNA was duplexed with the CRISPR-Cas9 tracrRNA (IDT, 1075927) by heating at 95°C for 5 min. This duplex was then complexed with recombinant Cas9 protein (Biotechne, 731-C3-005) by incubating at room temperature for 15 min. WM793 melanoma cells were detached with trypsin, washed in PBS and 5 x 10^5^ cells were electroporated in the presence of the crRNA/ tracrRNA.Cas9 complex using the Neon^TM^ Electroporation system (Thermofisher scientific, MPK5000) (4 pulses of 20ms at 1400V). Electroporated cells were plated with limit dilution to obtain clonal populations.

### siRNA transfection

To downregulate the genes of interest, we used RNA interference. 10^5^ A375, 2 × 10^5^ WM793, or 1.5 × 10^5^ WM852 cells were seeded in 6-well plate wells 24 h prior to transfection. Subsequently, cells were transfected with 25 nM or 50 nM siRNAs targeting specific genes: *CASP3* (Qiagen, 1027417, SI02654603; Horizon Discovery, L-004307-00-0005; Sigma-Aldrich, SASI_Hs01_00139105), *CASP7* (Sigma-Aldrich, SASI_Hs01_00128361), *CORO1B* (Sigma-Aldrich, SASI_Hs01_00010141, SASI_Hs01_00010142), and *SP1* (Sigma-Aldrich, SASI_Hs02_00363664). The negative control used was a non-targeting siRNA (Sigma-Aldrich, SIC001-10NMOL).

For protein and RNA analysis, cells were transfected for 48 h prior to analysis. For *in vitro* and *in vivo* cell phenotype analysis (motility assay, clonogenicity assay) and immunofluorescence experiments, cells were transfected for 24 h and then seeded *de novo* into the appropriate plates or injected (for zebrafish experiments) for subsequent experimentation.

### RNA sequencing and quantitative RT-PCR

Total RNA extraction was carried out using the Nucleospin RNA extraction kit (Macherey Nagel, 740955) according to the manufacturer’s protocol. For RNA sequencing (RNAseq) analysis, libraries were constructed using 600 ng of total RNA per sample with the TruSeq Stranded mRNA kit (Illumina), following the manufacturer’s guidelines. The process involved capturing PolyA mRNA using oligo-dT beads, synthesizing double-stranded cDNA, ligating adapters, amplifying libraries, and sequencing. Sequencing was performed using the NextSeq500 Illumina sequencer with 75 bp paired-end reads.

For quantitative real-time PCR (qRT-PCR) analysis of mRNA expression levels, cDNA synthesis was performed using the Sensifast cDNA synthesis kit (Bioline, BIO-65053). Gene-specific primers were designed using the Primer-BLAST tool available at the NCBI website (https://www.ncbi.nlm.nih.gov/tools/primer-blast/) and are detailed in Table S1. *GAPDH* was used as the reference gene for normalization. The qRT-PCR thermal cycling conditions included an initial polymerase activation step at 95°C for 2 min, followed by 40 cycles of denaturation at 95°C for 5 s and annealing/extension at 60°C/72°C for 30 s each.

### Western blot analysis

Protein extraction was carried out using RIPA buffer (Cell Signaling, 9806S), supplemented with protease inhibitor cocktail (Sigma-Aldrich, 4693116001) and phosphatase inhibitors (Sigma-Aldrich, P5726-1ML, P6044-1ML). Protein concentrations were determined using the Protein Assay Dye Reagent Concentrate (Bio-Rad, 5000006). Thirty micrograms of each protein sample were separated by SDS-PAGE and transferred onto a nitrocellulose membrane using the Transblot Turbo Transfer System (Bio-Rad, 1704150EDU). Non-specific binding sites on the membranes were blocked with 5% BSA in TBS-Tween 0.1% for 60 min. Next, the membranes were incubated overnight at 4°C with primary antibodies diluted at 1/1000 in 1% BSA TBS-Tween 0.1%. The primary antibodies used included CASP3 (Clinisciences, sc-56053; Cell Signaling, 9662S; Euromedex, GTX110543), CASP7 (Cell Signaling, 12827S), SP1 (Cell Signaling, 9389S), CORO1B (Thermo Fisher Scientific, CF501573), P-CORO1B (Euromedex, EC-CP2621), GFP (Origene, TP401), Myc-tag (Thermo Fisher Scientific, 13-2500), BAX (Cell Signaling, 2772S), BAK (Cell Signaling, 12105S), P-PKCα (Invitrogen, 44-962G), PKCα (Cell Signaling, 2056T), ARP2/ARP3 (Bioss Antibodies, bs-12524R), HSC70 (Clinisciences, sc-7298), Actin (Sigma-Aldrich, A3854), GAPDH (Sigma-Aldrich, CS207795), and vimentin (Sigma-Aldrich, CS207806). Following primary antibody incubation, membranes were washed three times in TBS-T. Subsequently, membranes were incubated with the appropriate secondary antibody coupled to an infrared fluorescent dye (Li-Cor Biosciences, 926-68070 and 926-32211). Membranes were washed three times and analyzed using the Li-Cor Odyssey Clx Infrared Fluorescent Western Scanning System (Li-Cor Biosciences).

For biotinylation experiments, detection of biotinylated proteins was performed following the blocking step. Streptavidin coupled to horseradish peroxidase was used at a concentration of 100 ng/mL (Sigma-Aldrich, OR03L-200UG) and incubated with the membranes for 30 minutes at room temperature with agitation. Subsequently, membranes were washed three times to remove unbound reagents. Biotinylated proteins were visualized using Clarity Western ECL blotting substrates (Bio-Rad, 1705060) and imaged using the ChemiDoc imager (Bio-Rad, 17001401).

### Immunofluorescence

Cells (5 x 10^4^ WM793 or WM852) were seeded onto coverslips in 24-well plates 24 h prior to staining, optionally coating the coverslips with Matrigel (Sigma-Aldrich, E6909-5ML) at a concentration of 100 µg/mL for 1 h for focal adhesion-associated protein analysis. After incubation, cells were fixed with 4% paraformaldehyde (PFA) for 5 min and washed three times with PBS. Permeabilization was achieved using 0.2% Triton X-100 in PBS for 10 min, followed by blocking of non-specific binding sites with 2% BSA in PBS. Next, cells were incubated with the following primary antibodies: Paxillin (1/400, BD Transduction Biosciences, 610052), P-CORO1B (1/100, Euromedex, EC-CP2621), CORO1B (1/100, Origene, CF501573), ARP2/3 (1/100, Bioss Antibodies, bs-12524R), and CASP3 (1/300, Euromedex, GTX110543), for 90 min at room temperature. After three washes with PBS, cells were incubated with appropriate secondary antibodies coupled to Alexa Fluor (1/300, Thermo Fisher Scientific, A21151, A21206, A31571) and, if required, with phalloidin-TRITC (1/300, Sigma-Aldrich, P1951), for 1 h at room temperature protected from light. Following three washes with PBS, cells were mounted with DAPI-containing VECTASHIELD® mounting medium (Eurobio Scientific, H-1200).

Alternatively, for the identification of biotinylated protein localization, after blocking non-specific binding sites, biotinylated proteins were stained with streptavidin-FITC (1/200, Biolegend, 405201) and phalloidin-TRITC (1/300, Sigma-Aldrich, P1951) for 30 min at room temperature protected from light. Slides were then washed and mounted as previously. Imaging was performed using a confocal Zeiss 980 microscope (Zeiss).

### Protein proximity analysis (PLA)

Cells (5 x 10^4^ WM793 or WM852) were seeded onto coverslips in 24-well plates and incubated for 24 h. Following incubation, cells were washed and fixed in 4% PFA for 5 min. After fixing, cells were washed three times with PBS. Protein proximity was analyzed using the Duolink® approach according to the manufacturer’s instructions (Sigma-Aldrich). Briefly, non-specific binding sites were blocked by incubating slides in Duolink® blocking buffer for 60 min at 37°C. Next, slides were incubated with primary antibodies: CORO1B (1/200, Origene, CF501573) and CASP3 (1/600, Euromedex, GTX110543), for 1 h at room temperature. After washing, cells were incubated with the secondary antibodies Duolink® In Situ PLA® Probe Anti-Rabbit PLUS (Sigma-Aldrich, DUO92002-100RXN) and Duolink® In Situ PLA® Probe Anti-Mouse MINUS (Sigma-Aldrich, DUO92004-100RXN) for 1 h at 37°C. Proximity of the secondary antibodies was then detected using Duolink® In Situ Detection Reagents Red (Sigma-Aldrich, DUO92008-100RXN), which includes a ligation and amplification step. Finally, images were acquired using a confocal Zeiss 980 microscope (Zeiss).

### Adhesion assay

To assess cell adhesion, 96-well ImageLock plates (Sartorius, 4379) were coated with Matrigel (Sigma-Aldrich, E609-10mL) at 100 µg/mL for 1 h at 37°C. Excess Matrigel was removed, and 10^3^ WM793 melanoma cells were seeded onto the remaining thin layer. Cell images were acquired at various time intervals using the IncuCyte ZOOM Imaging System (Sartorius), and cell adhesion was determined through cell shape analysis and scoring.

### Wound healing assay

For migration assays, 3 x 10^4^ WM852 or 5 x 10^4^ WM793 cells were seeded in a 96-well ImageLock plate (Sartorius, 4379) and allowed to form a monolayer for 24 h prior to the assay. Subsequently, a wound was created in the cell monolayer using the WoundMaker (Sartorius, 4563) as per the manufacturer’s instructions. Cell migration to close the wound was assessed by time-lapse microscopy using the IncuCyte ZOOM imaging system.

### Invasion assay

Invasion capacity was evaluated using a 96-well ImageLock plate (Sartorius, 4379) coated with Matrigel (Sigma-Aldrich, E609-10mL) at 100 µg/mL for 1 h at 37°C. Excess Matrigel was removed, and 3 x 10^4^ WM852 or 5 x 10^4^ WM793 cells were seeded 24 h prior to the assay. A wound was created in the cell monolayer using the WoundMaker, followed by the addition of a new layer of Matrigel (800 µg/mL) that was allowed to polymerize for 1 h at 37°C. Cell invasion was evaluated using time-lapse microscopy with the IncuCyte ZOOM system.

### Chemotaxis assay

The chemotactic capacity of melanoma cells was determined using xCELLigence technology. 3 x10^4^ melanoma cells (WM793 or WM852) were seeded onto the upper chamber of a CIM-Plate 16 (ACEA Biosciences, 05665817001), while the lower chamber was filled with medium supplemented with 20% fetal bovine serum. Melanoma cell migration was monitored over time using the xCELLigence RTCA DP instrument (ACEA Biosciences).

### Analytical flow cytometry

2 x 10^5^ WM793 cells or 1.5 x 10^5^ WM852 cells stably expressing the VC3ai caspase reporter construct were seeded in 6-well plates 24 h before treatment. The cells were then treated with apoptosis inducers for 24 h, and VC3ai cleavage by caspase-3/7 was assessed based on fluorescence emission at 530 nm using a BD LSR Fortessa™ Cell Analyzer (BD Biosciences).

### Cytoskeleton protein isolation

Cytoskeleton fraction characterization was performed using the ProteoExtract® Cytoskeleton Enrichment and Isolation Kit (Sigma-Aldrich, 17-10195) according to the manufacturer’s instructions.

### Holotomographic microscopy

Control and CASP3-depleted WM793 cells (10^4^) were seeded onto Fluorodishes (Ibidi GmbH, Gräfeling, Germany). Holotomographic microscopy was conducted using the 3D Cell-Explorer Fluo (Nanolive, Ecublens, Switzerland) with a ×60 air objective at a wavelength of λ = 520 nm. Live-cell imaging was performed under physiological conditions using a top-stage incubator (Oko-lab, Pozzuoli, Italy) to maintain a constant temperature of 37°C, 5% CO_2_ level, and air humidity saturation. Refractive index maps were generated every 5 min for 1 h, and images were processed using STEVE software.

### Zebrafish tail veins metastasis model

Prior to injection, melanoma cells (9 x 10^5^) were suspended in serum-free medium and stained with lipophilic dyes DiO or DiD for 20 min at 37°C (Thermo Fisher Scientific, V22889). After staining, cells were washed and resuspended in 30 µL of PBS. For zebrafish xenotransplantation, zebrafish embryos at 48 h post-fertilization (hpf) were dechorionated and anesthetized using tricaine (Sigma-Aldrich, E10521). Approximately 300 labelled human cells in 20 nL of suspension were injected into the perivitelline cavity of each embryo. The injected embryos were then incubated at 30°C for 24 - 48 h and allowed to recover in the presence of N-phenylthiourea (Sigma-Aldrich, P7629) to inhibit melanin synthesis. For imaging and metastasis assessment, zebrafish embryos were anesthetized with tricaine and imaged using an Axio Observer Zeiss microscope (Zeiss).

### Zebrafish embryo microinjection

Tg(mitfa:*BRAF^V600E^*), *tp53*^−/−^, *mitfa*^−/−^ zebrafish were bred, and embryos were collected for microinjection. Embryos at the one-cell stage were injected with MiniCoopR vectors for the expression of CASP3 or the control gene GFP. Each embryo received 25 pg of DNA constructs and 25 pg of Tol2 mRNA. Post-injection, embryos were raised in E3 medium at 28.5°C. Melanocyte rescue was assessed 4 days post-fertilization. Tumor development was monitored from 11 to 20 weeks post-injection. Pigmented tumors were randomly collected from adult male and female fish, aged 3 to 6 months, for transplantation studies.

MiniCoopR is a DNA plasmid that can be stably integrated into the zebrafish genome via Tol2-mediated transposition and enables gene expression under the control of the melanocyte-specific mitf promoter.

#### In vivo tumour spreading assay

Primary zebrafish melanoma cells were transplanted into irradiated adult *casper* recipients following a previously established protocol. Zebrafish aged 3 to 6 months were subjected to a total of 20 Gy irradiation over two days. A suspension of 300,000 melanoma cells in 3 μl HBSS was then subcutaneously injected into their dorsal cavity using a 10 μl Hamilton syringe fitted with a 32-gauge bevel-tipped needle. Each primary tumor was transplanted into 8 to 12 secondary recipients. Pictures of transplanted fish were taken 21 days post-transplantation to evaluate spreading.

Experiments on zebrafish were performed at the PRECI Aquatics Facility that has been approved by the Department of Population Protection – Prefecture du Rhône (under number C69387 0602). All procedures involving the use of animals are submitted for review and approval by the local Animal Ethics Evaluation Committee (CECCAPP) and for validation by the French Ministry of Higher Education, Research and Innovation. The protocol for the generation of genetically-modified primary melanoma in adult zebrafish and the transplantation of zebrafish melanoma cells into adult transparent recipients has been approved under number APAFIS#34134-20210620122832.

### Coimmunoprecipitation using GFP-Trap ®

CASP3-GFP interacting partners were characterized using GFP-Trap® Magnetic Agarose (20 reactions, Euromedex, CR-gtma-20) following the manufacturer’s instructions. Briefly, melanoma cell lines expressing GFP or CASP3-GFP (WM793 or WM852) at 80% confluency in 150-mm culture plates were lysed in NP40 buffer (10 mM Tris/HCl pH 7.4, 150 mM NaCl, 0.5 mM EDTA, 0.5% NP40, and protease/phosphatase inhibitors) on ice for 30 min, followed by centrifugation at 10,000 × g for 5 min. The resulting cell lysates were incubated with 25 µL of GFP-Trap® Magnetic Agarose beads overnight at 4°C. Beads were then washed three times with wash buffer (10 mM Tris/HCl pH 7.5, 150 mM NaCl, 0.05% NP40). 10% of the beads were resuspended in Laemmli buffer for Western blot analysis, while the remaining 90% were resuspended in 50 mM ammonium bicarbonate buffer for subsequent mass spectrometry analysis.

For the validation of specific interactants, GFP and CASP3-GFP melanoma cell lines (WM793 and WM852) were resuspended in PBS supplemented with protease and phosphatase inhibitors and mechanically lysed by passing cells several times through a 29G syringe. Cell lysates were then incubated with 25 µL of GFP-Trap® Magnetic Agarose beads overnight at 4°C. Beads were washed three times with PBS. GFP and CASP3-GFP protein complexes were eluted using Laemmli buffer and samples were denatured at 95°C for further investigation by Western blot analysis.

### Proximity biotinylation assay

On the first day, 10^6^ WM793 cells or 7.5 × 10^5^ WM852 cells stably overexpressing either pITR-mycBioID2, pITR-mycBioID2-CASP3, or pITR-CASP3-mycBioID2 were seeded onto a 100-mm plate. After 24 h, expression of mycBioID2, mycBioID2-CASP3, and CASP-mycBioID2 was induced by treatment with 1 µg/mL doxycycline, followed by overnight treatment with 50 mM biotin. On the third day, cells were washed with PBS and scraped into lysis buffer (0.2% SDS, 2% Triton X-100, 50 mM Tris/HCl pH 7.4, 500 mM NaCl, and protease/phosphatase inhibitors). The samples were sonicated at 30% amplitude for 30 s and then centrifuged at 10,000 × g for 10 min at 4°C. The resulting cell lysates were incubated with 50 µL of Streptavidin Dynabeads™ MyOne™ C1 beads (Thermo Fisher Scientific, 65001) overnight with rotation at 4°C. Beads were washed twice with 2% SDS, 50 mM Tris/HCl pH 7.4, followed by one wash each with 0.1% deoxycholic acid, 1 mM EDTA, 500 mM NaCl, 50 mM HEPES pH 7.5, and 0.5% deoxycholic acid, 1 mM EDTA, 250 mM LiCl, 10 mM Tris/Cl pH 7.4. 10% of the beads were resuspended in Laemmli buffer for Western blot validation, while the remaining 90% were resuspended in 50 mM ammonium bicarbonate buffer for subsequent mass spectrometry analysis.

### Caspase-3 fluorometric assay

Caspase-3 activity was assessed utilizing the Caspase 3/CPP32 Fluorometric assay kit, following the guidelines provided by the manufacturer (BioVision, K105).

### Plasmid construct

The pEGFP-N3-CASP3 fusion construct was generated by inserting the *CASP3* open reading frame into the pEGFP-N3 vector, after linearization with XhoI and SalI restriction enzymes (NEB, R0146S and R3138T). The *CASP3* gene was amplified from the *CASP3* coding plasmid (Addgene, 11813) using specific primers (detailed in Table S1). The PCR product was digested with XhoI and SalI, and the resulting fragment was purified using the NucleoSpin Gel and PCR Clean-up kit according to the manufacturer’s instructions (Macherey-Nagel, 740609.250). The purified *CASP3* gene fragment was ligated into the linearized pEGFP-N3 vector using the quick ligation^TM^ kit according to the manufacturer’s instructions (NEB, B2200S). The resulting mixture was transformed into competent DH5α bacteria (NEB, C2987I), and the plasmid was amplified and purified using the Nucleobond Xtra Midi kit (Macherey-Nagel, 740410.50).

Additionally, guide RNAs for targeting *CASP3* gene were inserted into the pLenti CRISPR V2 (addgene, 83480) allowing concomitant expression of Cas9. For this, the pLenti CRISPR V2 was digested for 2 h at 50°C with *Bsmb1* (NEB, R0580S) and the resulting digested fragment was purified using the NucleoSpin Gel and PCR Clean-up kit (Macherey-Nagel, 740609.250). Guide RNAs were designed using the CRISPOR tool (http://crispor.gi.ucsc.edu/) and are listed in Table S1. 100µM gDNAs were phosphorylated and annealed with T4 PNK (NEB, M0201S) according to the manufacturer’s instructions, and the resulting oligonucleotides were diluted 200X. Digested plasmid and oligonucleotides were ligated as previously described and transformed into Stbl3 bacteria.

Plasmids for the inducible expression of CASP3-BioID2 and BioID2-CASP3 were developed in two successive steps. First, CASP3 open reading frame and BioID2 either in N-terminus or in C-terminus of CASP3 were inserted into a pcDNA3 vector using the In-Fusion® HD Cloning kit (Takara, 639649) according to the manufacturer’s instructions. For this, the pcDNA3 vector was linearized using BamHI (Neb, R3136S) and EcoRI (NEB, R3101T). CASP3 was amplified from pcDNA3-Casp3-myc (Addgene, 11813) and mycBioID2 was amplified from mycBioID2-pBABE-puro (Addgene, 80900) using primers listed in Table S1 (designed using the Takara in-Fusion Cloning Design Primer tool (https://takara.teselagen.com/#/DesignPage)). Secondly, mycBioID2, BioID2-CASP3 and CASP3-BioID2 were inserted into the pITR2 vector. For this, mycBioID2 was amplified from mycBioID2-pBABE-puro (Addgene, 80900), CASP3-mycBioID2 and mycBioID2 – CASP3 were amplified from the previous developed pcDNA3 vectors using primers listed in table S1. PCR fragments and the linearized pITR2 vector were digested with SfiI (NEB, R0123S), purified, ligated and transformed into Stbl3 bacteria as previously described.

### Mass spectrometry analysis

#### Sample preparation

IP streptavidin beads were prepared with the iST kit (ref. P.O.0001, PreOmics) according to the manufacturer protocol for Magnetic Immunoprecipitation Samples. Briefly, proteins were denaturated, reduced and alkylated (10min, 60°C, 1000 rpm), then digested with a mixture of endoproteinase Lys-C/trypsin (3h, 37°C, 500 rpm). After a cleaning step, peptides were dried, suspended in Formic Acid (FA) 0.1% and dosed with the quantitative fluorometric peptide assay (ref. 23290, Thermo Scientific).

#### LC-MSMS analysis

100ng of each sample were analyzed on the Exploris 480 mass spectrometer coupled with a Vanquish NEO nanoLC system (ThermoFisher Scientific). Peptides samples were loaded on a C18 Acclaim PepMap100 trap-column 300 µm ID x 5 mm, 5 µm, 100Å (ThermoFisher Scientific) and separated on a C18 Acclaim Pepmap100 nano-column (ThermoFisher Scientific) with a 30 minutes linear gradient from 3% to 35% buffer B (A: 0.1% FA in H_2_O, B: 0.1% FA in ACN/H_2_O (80/20)) followed by a washing and equilibration steps for a total duration of 40 minutes. The flow rate was 300 nL/min and the oven temperature was kept constant at 45°C. Peptides were analysed with a DDA 1s HCD method: MS data were acquired in a data dependent strategy (DDA) selecting the fragmentation events based on the most abundant precursor ions in a 1s survey scan (350-1400 Th). Resolutions of the survey and MS/MS scans were respectively set at 120,000 and 15,000 at m/z 200 Th. The Ion Target Values for the survey and the MS/MS scans in the Orbitrap were set to 3E6 (300%) and 1E5 (100%) respectively and the maximum injection time was set to 50 ms for MS scan and 22 ms for MS/MS scan. Parameters for acquiring HCD MS/MS spectra were as follows : collision energy = 30 and isolation window = 4 m/z. The precursors with unknown charge state, charge state of 1 and 8 or greater than 8 were excluded. Peptides selected for MS/MS acquisition were then placed on an exclusion list for 30 s using the dynamic exclusion mode to limit duplicate spectra.

#### Data analysis

Raw data were processed with Proteome Discoverer 3.1 through the SEQUEST HT and CHYMERIS search engines against the *homo sapiens* database (Uniprot, Release June 2023, 20348 sequences) and a database of common contaminants. Precursor mass tolerance was set at 10 ppm and fragment mass tolerance was set at 0.02 Da, and up to 2 missed cleavages were allowed. Oxidation (M), acetylation (Protein N-terminus) were set as variable modification and Carbamidomethylation (C) as fixed modification. Validation of identified peptides and proteins was done using a target decoy approach with a false discovery rate positive (FDR < 1%) via Percolator. Protein quantification was done by a Label Free Quantification (LFQ) approach. LFQ abundance values were normalized to the total peptide amount and ratios of quantitation were calculated in a pairwise way. Statistical validation was based on t-test and a protein is considered differentially expressed between two conditions if the fold change is > 2 or < 0.5 and have a p-value < 0.05.

### Bioinformatics analysis

Illumina sequencing was performed on RNA extracted from triplicates of each condition. Standard Illumina bioinformatics analysis were used to generate fastq files, followed by quality assessment [MultiQC v1.7 https://multiqc.info/]. ‘Rsubread’ v1.34.6 was used for mapping to the hg38 genome and creating a matrix of RNA-Seq counts. Next, a DGElist object was created with the ‘edgeR’ package v3.26.7 [https://doi.org/10.1093/bioinformatics/btp616]. After normalization for composition bias, genewise exact tests were computed for differences in the means between groups, and differentially expressed genes (DEGs) were extracted based on an FDR-adjusted p value < 0.05 and a minimum absolute fold change of 4. The RNA-seq data was submitted to the GEO repository (GSE270452).

Caspase 3 expression in normal tissues was determined using the ENSG00000164305.18 dataset from The Genotype-Tissue Expression website (https://gtexportal.org/home/gene/ENSG00000164305). Caspase expression in 39 melanoma cell lines was assessed in the Cancer Cell Line Encyclopedia (CCLE) (https://www.ebi.ac.uk/gxa/experiments/E-MTAB-2770/Results). Mutations of *CASP3* were addressed using the Catalogue of Somatic Mutations in Cancer (COSMIC) (https://cancer.sanger.ac.uk/cosmic). Association of CASP3 expression with metastasis formation in melanoma (SKCM) was analyzed using The Cancer Genome Atlas (TCGA) (https://portal.gdc.cancer.gov/). The correlation between *CASP3* and *SP1* expression was assessed using the GEPIA2 online tool with TCGA-SKCM data (http://gepia2.cancer-pku.cn/#index). Concerning RNA sequencing data, a siCASP3 upregulation (UP) signature was designed by selecting the 50 most upregulated genes following knockdown of CASP3. Gene Set Variation Analysis (GSVA) was utilized to calculate the score for each patient from TCGA-SKCM, and Kaplan-Meier analysis was performed to compare patients expressing a high or low score (according to the median). Additionally, non-supervised clustering was done with genes involved in lamellipodia assembly (GOBP_LAMELLIPODIUM_ASSEMBLY) showing that this signature is sufficient to separate mRNA from control cells and mRNA from cells transfected with CASP3 siRNA.

### Analysis of caspase-3 interactome

New potential interactants were identified through a process of filtering the protein list using 139 previously described interaction datasets, which were obtained from the OmniPath server on June 20, 2023 with the OmnipathR package (version 3.4.7) ^54^. Additionally, potential C3 substrates identified using degradase (V1, downloaded 2022/09/29) were excluded from further analysis ^55^.

Furthermore, InterPro protein domains annotations were retrieved from the OmniPath server. To streamline the analysis, domain summarization was conducted by considering only one occurrence of each domain.

Overrepresentation of Gene Ontology biological processes was performed on filtered protein list utilizing clusterProfiler package (version 4.4.4) ^56^.

**Supplementary Table 1.**
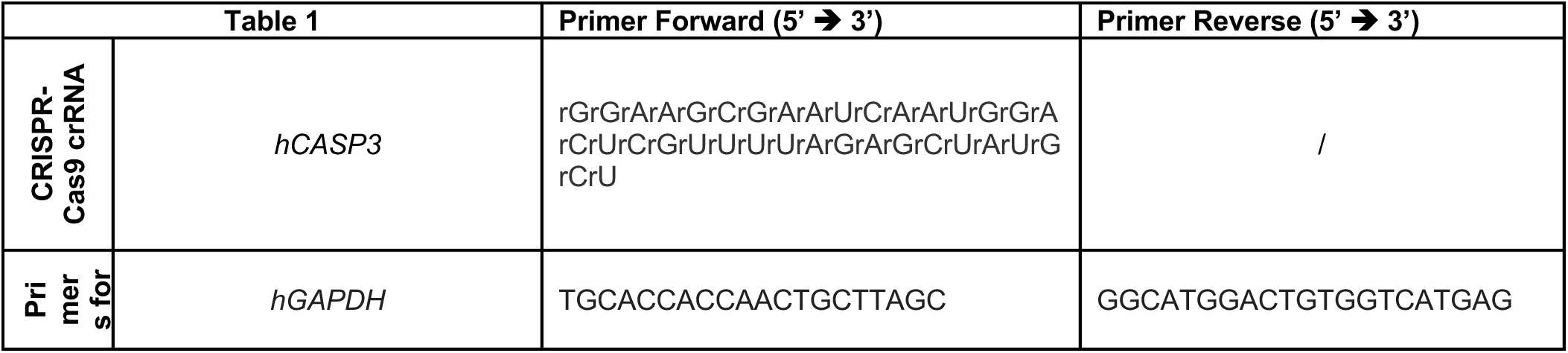
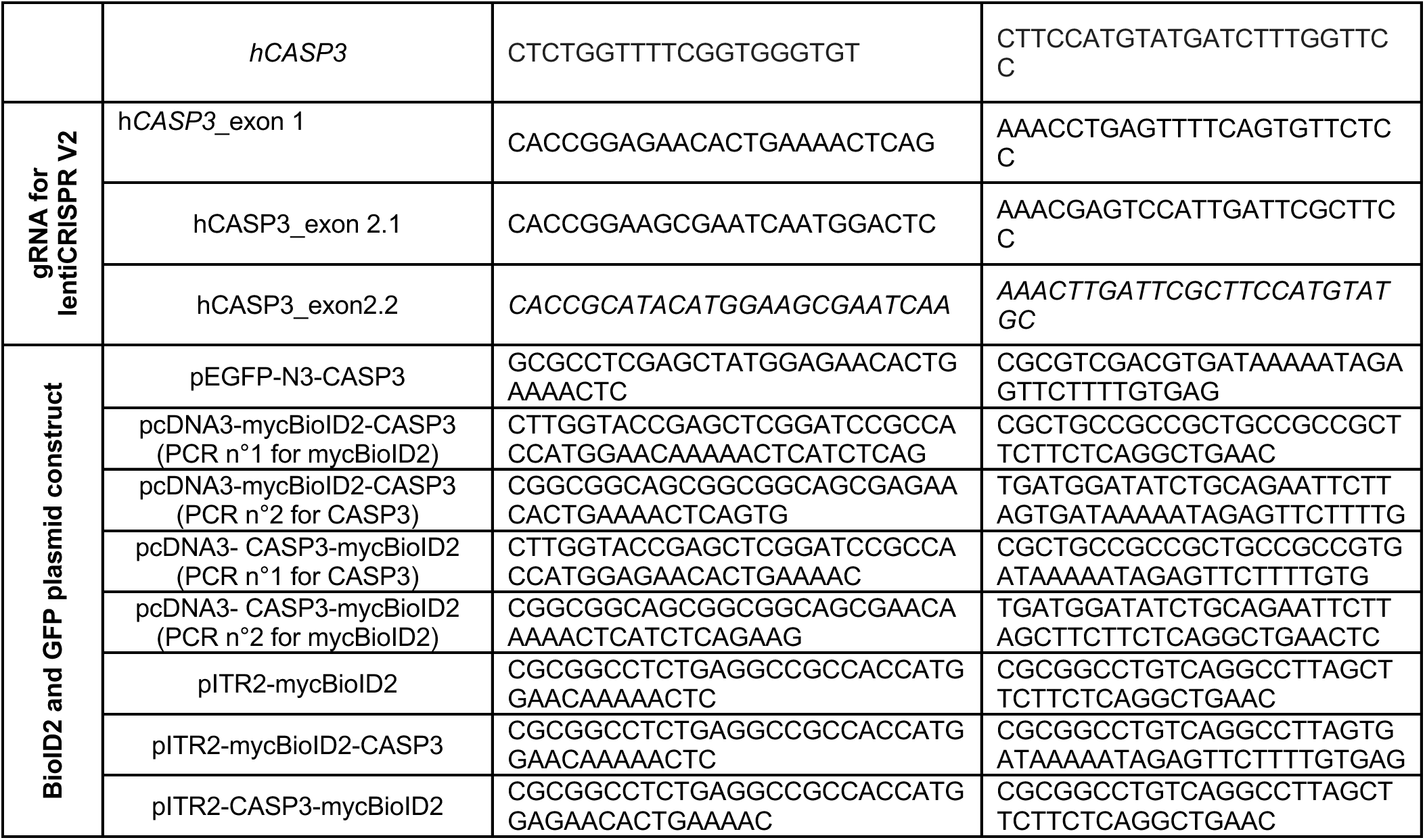
Primer List.

---

**Supplementary Figure 1.**
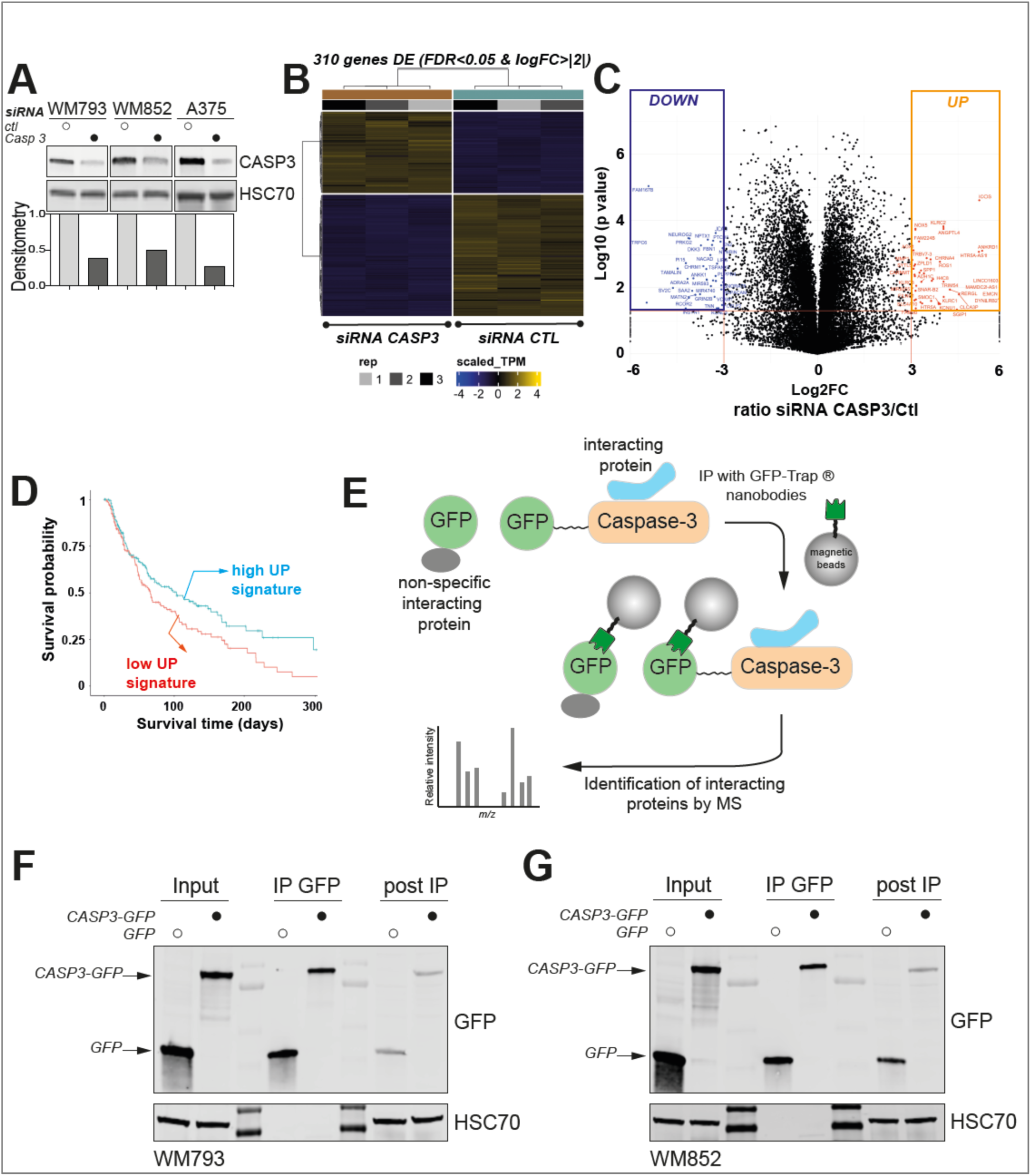
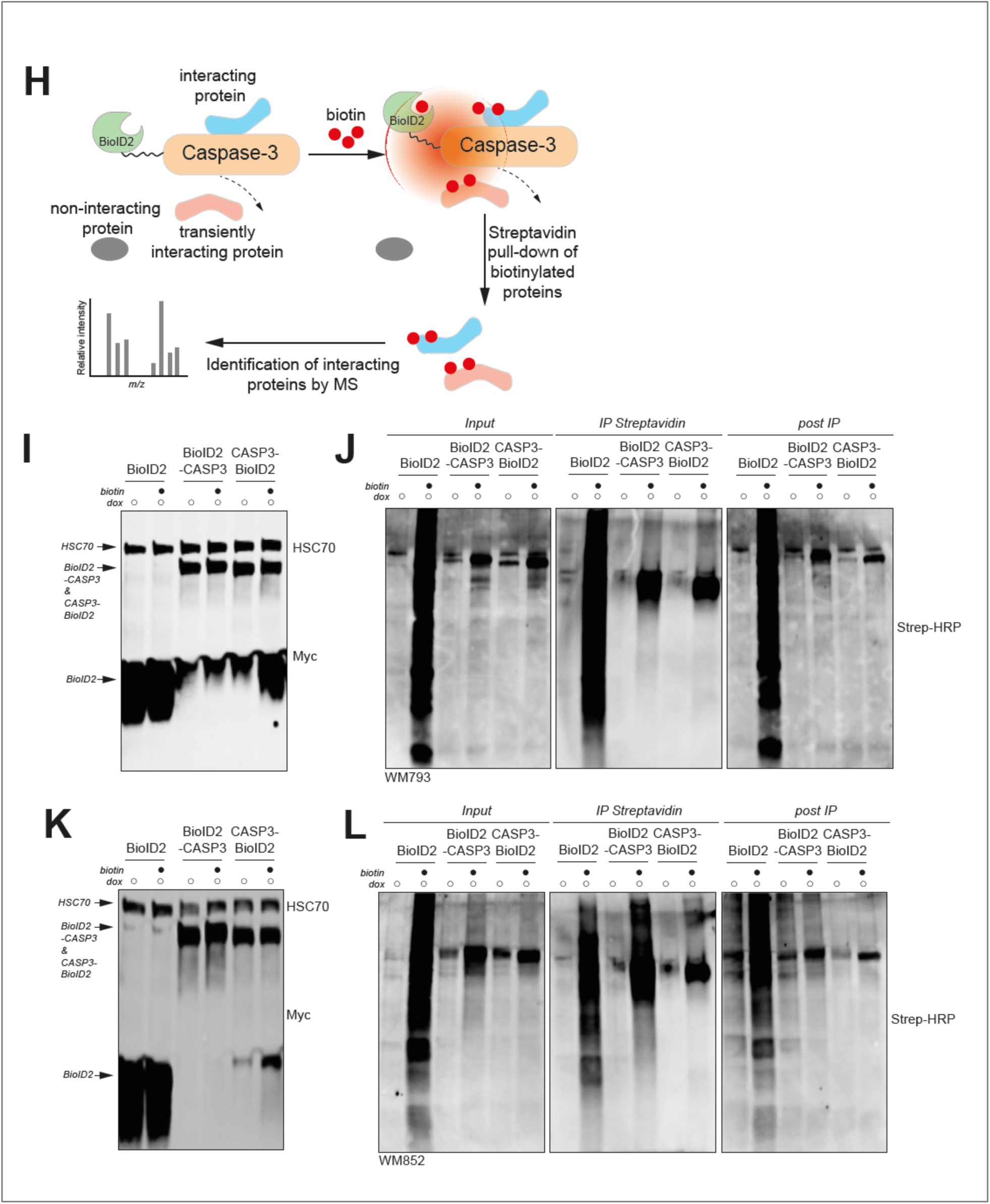
Related to Fig.1. **(A)** Upper panel: Western blot analysis of CASP3 in WM793, WM852 and A375 cells transfected with *CASP3*-targeting siRNA. HSC70 serves as a loading control. Lower panel: Densitometry analysis of CASP3 expression, relative to HSC70 expression. **(B)** Clustering of the most differentially-expressed genes (DEGs) between parental and CASP3-deficient WM793 cells, identifying a DEG signature constituted of 310 genes (FDR<0.05 & logFC>|2|). **(C)** Volcano plot illustrating the two gene signatures constituted of DEGs that are the most significantly upregulated or downregulated following CASP3-depletion in WM793. **(D)** Analysis of the overall survival in patients with melanoma from the TCGA dataset displaying a high or low (relative to the median) GSVA score for the signature of genes increased when caspase-3 is knocked-down (50 genes the most upregulated upon C3 knocked-down) (log rank test; p-value = 0.015). **(E)** The protocol used for identifying interaction protein partners for CASP3 using immunoprecipitation of CASP3-GFP with GFP Traps^®^, which are ready-to-use pull-down reagent consisting of an anti-GFP nanobody coupled to agarose beads. **(F-G)** Validation by immunoblotting of GFP and CASP3-GFP immunoprecipitation using GFP Traps^®^ in GFP- and CASP3-GFP-expressing WM793 (**F**) and WM852 cells (**G**). **(H)** Summary of key steps of the proximity labelling protocol for identifying protein interacting partners of CASP3 fused with BioID2 through the immunoprecipitation of biotinylated proteins with streptavidin beads, following biotin treatment. **(I-J)** Validation of myc tagged-BioID2, -BioID2-CASP3 and -CASP3-BioID2 conditional overexpression following doxycycline treatment (1 µg/mL for 24 h) using anti-Myc antibody in WM793 cells (**I**). Analysis of biotinylated proteins following biotin treatment (50 µM, 12 h) of BioID2, BioID2-CASP3 and CASP3-BioID2-expressing WM793 cells (**J**). (**K-L**) Same as in I-J, for WM852 cells.

**Supplementary Figure 2.**
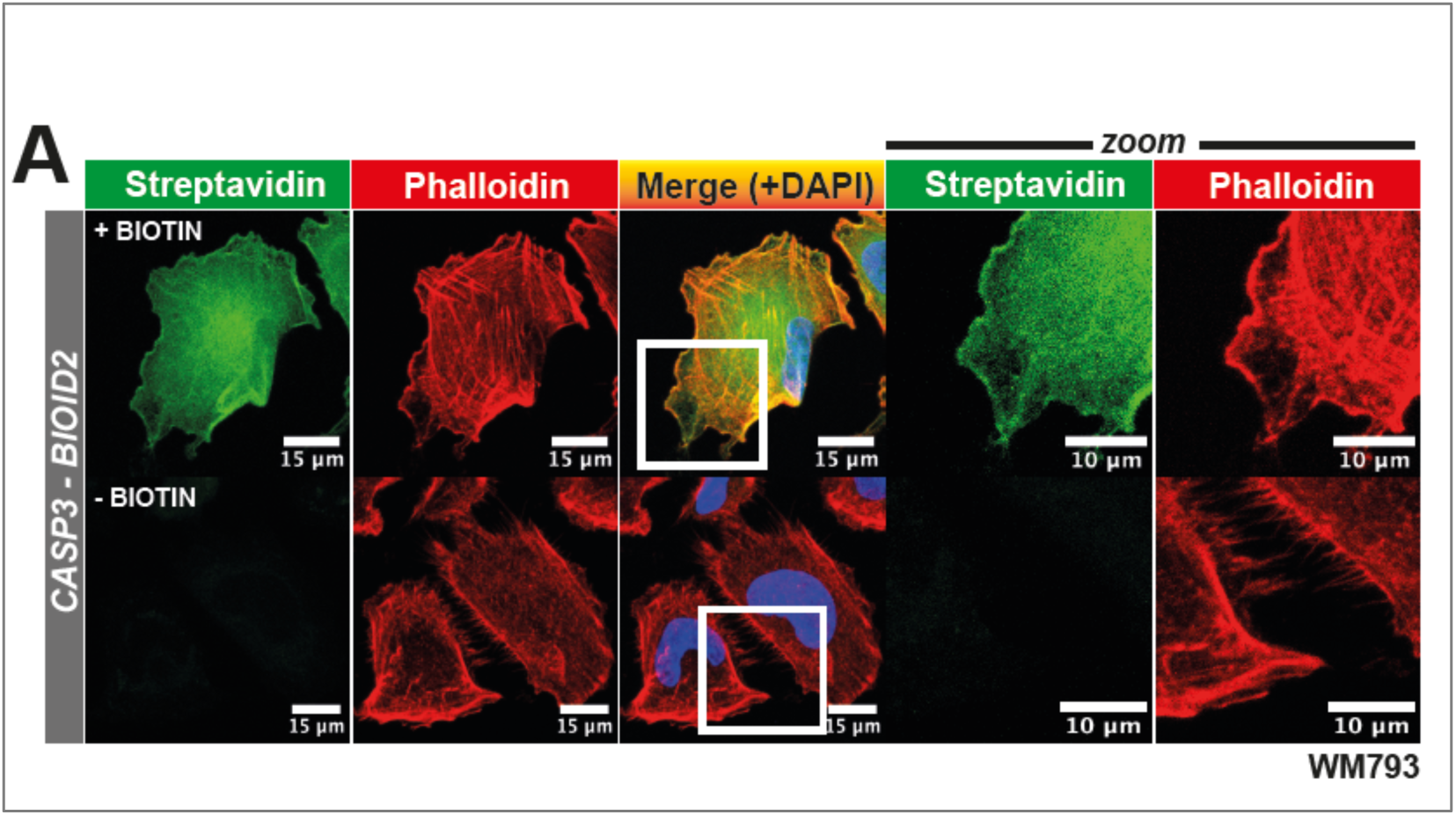
Related to Fig.2. **(A)** Biotinylated protein immunostaining using Streptavidin-FITC in WM793 overexpressing CASP3-BioID2.

**Supplementary Figure 3.**
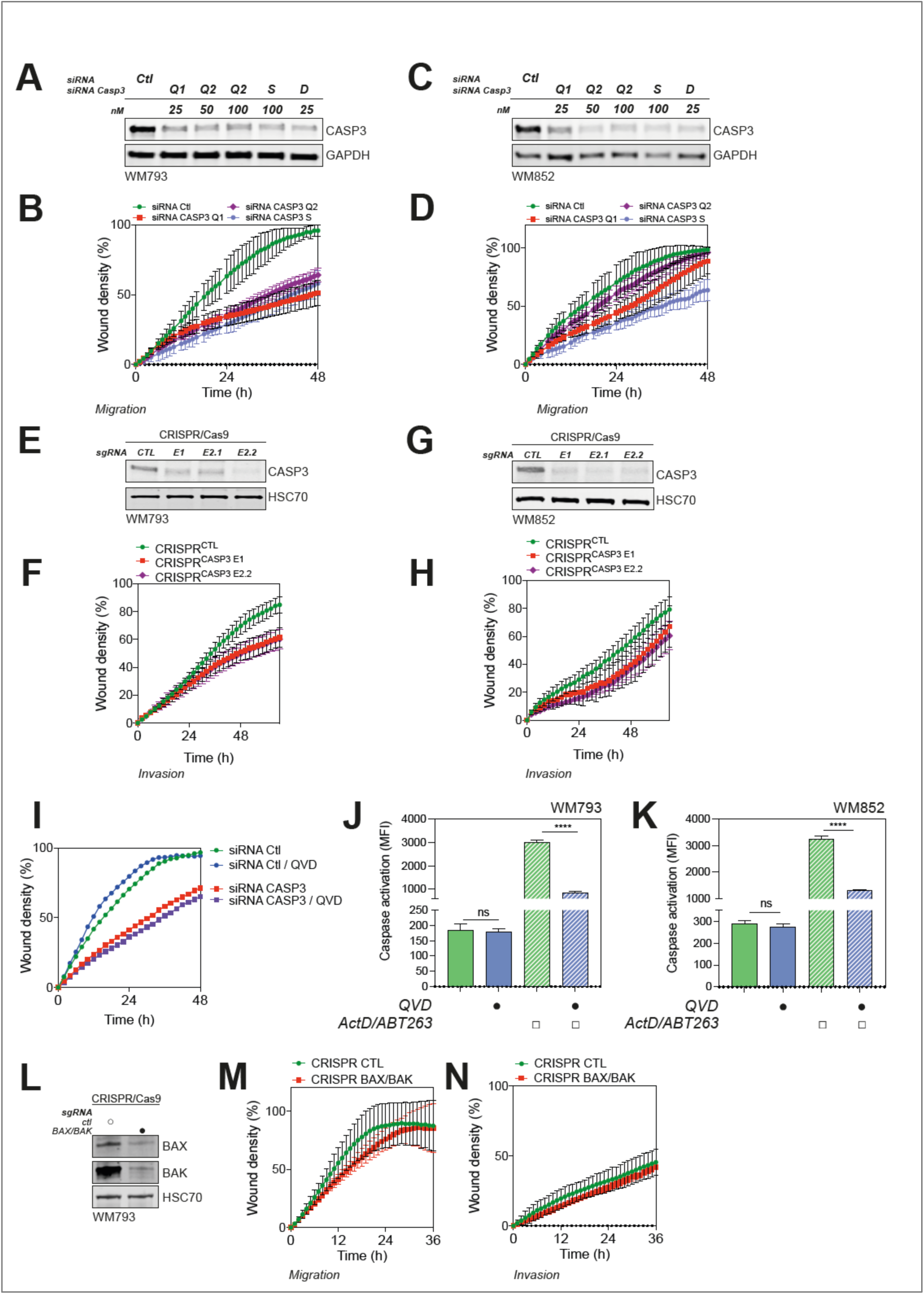
Related to Fig.3. **(A)** Immunoblotting analysis of CASP3 expression in WM793 transfected for 48 h with various CASP3 siRNA (Q1: Qiagen n°1; Q2: Qiagen n°2; S: Sigma-Aldrich; D: Dharmacon) at the indicated concentration. GAPDH serves as a loading control. **(B)** Quantification of the migration potential of parental and CASP3-deficient WM793 cells using various siRNAs, through wound area measurement (data represent mean with SD of a representative experiment). **(C)** Analysis by immunoblotting of CASP3 expression in WM852 transfected for 48 h with various CASP3 siRNA (Q1: Qiagen n°1; Q2: Qiagen n°2; S: Sigma-Aldrich; D: Dharmacon) at the indicated concentration. GAPDH serves as a loading control. **(D)** Quantification of the migration potential of control and CASP3-deficient WM852 cells using various siRNAs through wound area measurement (data represent mean with SD of a representative experiment). **(E)** Analysis by immunoblotting of CASP3 expression in CRISPR/Cas9-edited WM793 cells, electroporated with either control or *CASP3*-targetting sgRNAs. HSC70 serves as a loading control. **(F)** Quantification of the invasion potential of CRISPR/Cas9 control and *CASP3*-targetted WM793 cells, through wound area measurement (data represent mean with SD of a representative experiment). **(G-H)** Same as in E-F, for WM852 cells. **(I)** Quantification of the migration potential of control and CASP3-deficient WM793 cells treated with the pan-caspase inhibitor qVD-OPh (10µM) through wound area measurement (data represent mean with SD of a representative experiment). **(J-K)** Measurement of caspase-3/7 activation in WM793 (**J**) and WM852 cells (**K**) expressing the caspase reporter VC3AI, and treated with actinomycin D (ActD, 1 µM), ABT-263 (5 µM) and qVD-OPh (10 µM) for 24 h. **(L)** Analysis by immunoblotting of BAX and BAK expression in *BAX*/*BAK1* double knock-out CRISPR/Cas9-edited WM793 cells. HSC70 serves as a loading control. **(M-N)** Quantification of the migration (**M**) and invasion (**N**) potential of CRISPR/Cas9 control and and *BAX*/*BAK1* double knock-out WM793 cells (data represent mean with SD of a representative experiment).

**Supplementary Figure 4.**
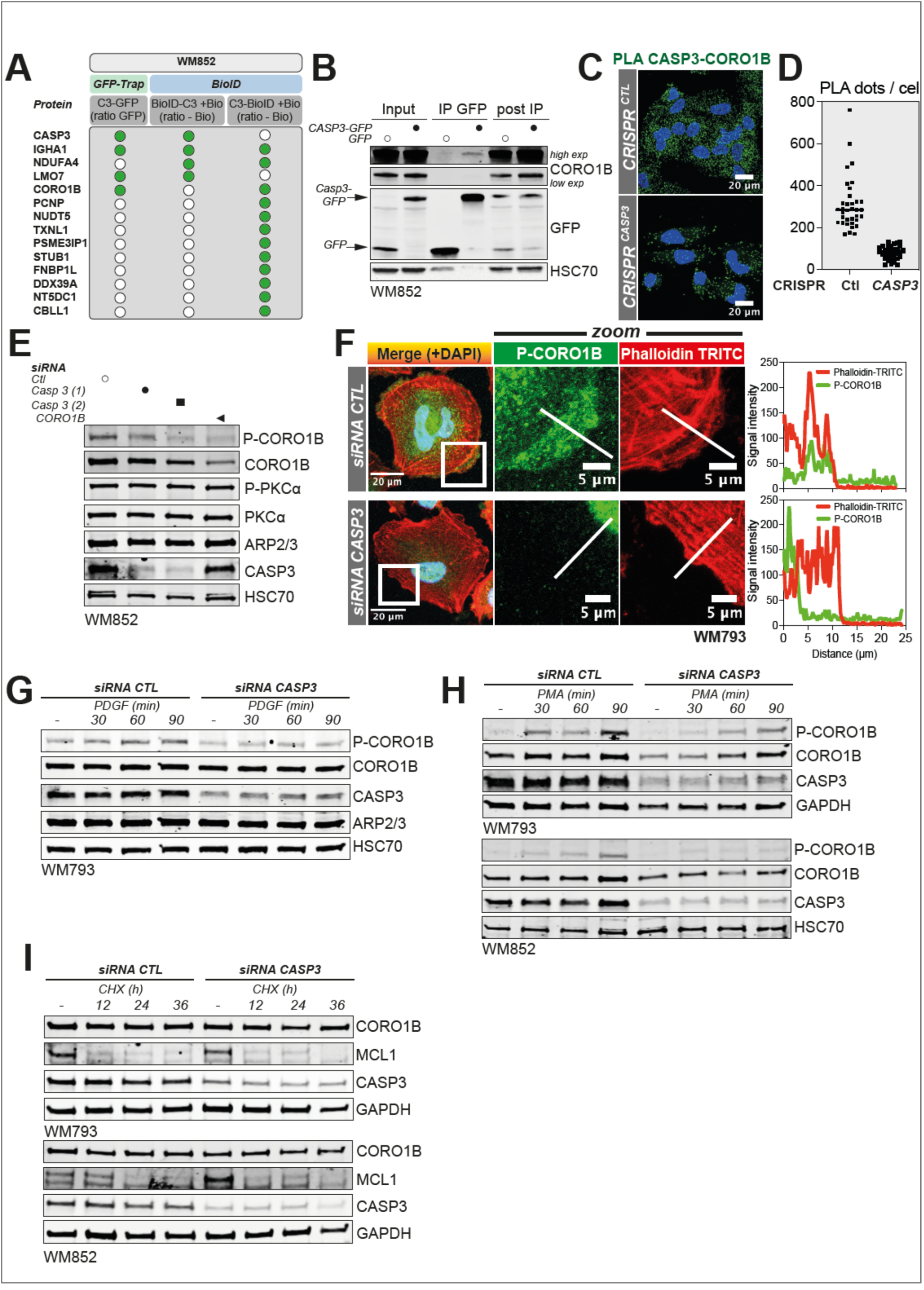
Related to Fig.5. **(A)** Summary table with the most frequent CASP3-interacting proteins identified in WM852 cells by CASP3-GFP pulldown and proximity labelling through biotinylation of neighboring proteins in cells expressing BioID2-fused CASP3. **(B)** Analysis of CASP3-GFP and CORO1B interaction after immunoprecipitation (IP) of GFP protein complexes in GFP- or CASP3-GFP in WM852 melanoma cells. **(C)** Analysis of the proximity between endogenous CASP3 and CORO1B proteins in CRISPR/Cas9 control and *CASP3*-targetted WM793 cells, using a Proximity Ligation Assay (PLA). **(D)** Quantification of PLA dots in in CRISPR/Cas9 control (n = 35 cells, in a representative experiment) and CASP3-targeted (n = 33 cells) WM793 cells. **(E)** Analysis by immunoblotting of P-CORO1B, CORO1B, P-PKCα, PKCα, ARP2/3, CASP3 in WM852 transfected with two different siRNA for CASP3 and CORO1B. HSC70 serves as a loading control. **(F)** Left panel: Analysis of P-CORO1B and F-actin localization by immunofluorescence in parental and CASP3-deficient WM793 cells. Right panel: Signal intensity measurement of P-CORO1B and F-actin in the indicated corresponding line in control and CASP3-deficient WM793 in the left panel. **(G)** Analysis by immunoblotting of P-CORO1B, CORO1B, ARP2/3 and CASP3 in parental and CASP3-deficient WM852 cells that were serum-starved for 24 h and treated with PDGF (20 ng/mL) for the indicated time. HSC70 serves as a loading control. **(H)** Analysis by immunoblotting of P-CORO1B, CORO1B, CASP3 in parental and CASP3-deficient WM793 and WM852 cells treated with PMA (100 nM) for the indicated time. HSC70 serves as a loading control. **(I)** Analysis by immunoblotting of CORO1B, CASP3 and MCL1 in control and CASP3-deficient WM793 and WM852 cells treated with cycloheximide (50 µg/mL) for the indicated time. GAPDH serves as a loading control.

**Supplementary Figure 5.**
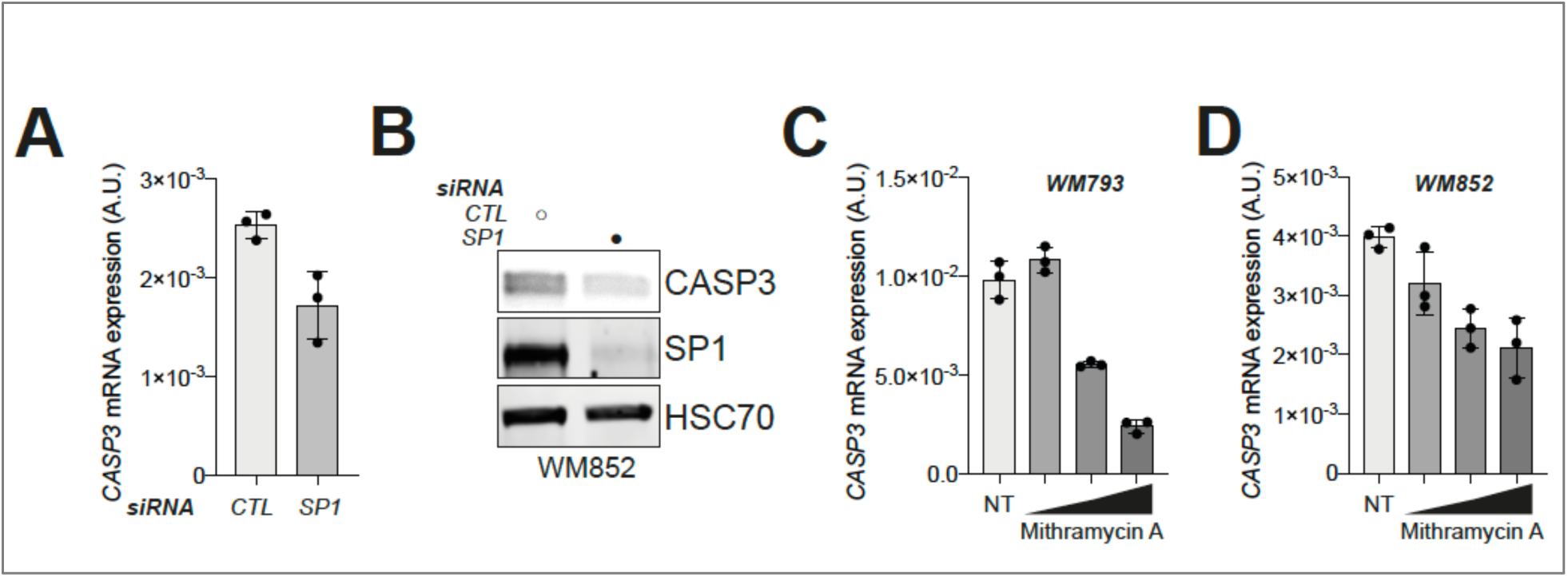
Related to Fig.6. **(A)** *CASP3* mRNA expression in control and SP1-deficient WM852 cells relative to *GAPDH* (Arbitrary Unit). **(B)** Western blot analysis of CASP3 and SP1 in control and SP1-deficient WM852 cells. HSC70 serves as a loading control. **(C-D)** *CASP3* mRNA expression in WM793 (**C**) and WM852 cells (**D**) treated with increasing doses of mithramycin A (100 nM, 200 nM and 300 nM for 24 h) relative to *GAPDH*.

